# Time-dependent material properties of ageing biomolecular condensates from different viscoelasticity measurements in molecular dynamics simulations

**DOI:** 10.1101/2022.12.07.519428

**Authors:** Andrés R. Tejedor, Rosana Collepardo-Guevara, Jorge Ramírez, Jorge R. Espinosa

## Abstract

Biomolecular condensates are important contributors to the internal organization of the cell material. While initially described as liquid-like droplets, the term biomolecular condensates is now used to describe a diversity of condensed phase assemblies with material properties extending from low to high viscous liquids, gels, and even glasses. Because the material properties of condensates are determined by the intrinsic behaviour of their molecules, characterising such properties is integral to rationalising the molecular mechanisms that dictate their functions and roles in health and disease. Here, we apply and compare three distinct computational methods to measure the viscoelasticity of biomolecular condensates in molecular simulations. These methods are the shear stress relaxation modulus integration (SSRMI), the oscillatory shear (OS) technique, and the bead tracking (BT) method. We find that, although all of these methods provide consistent results for the viscosity of the condensates, the SSRMI and OS techniques outperform the BT method in terms of computational efficiency and statistical uncertainty. We, thus, apply the SSRMI and OS techniques for a set of 12 different protein/RNA systems using a sequence-dependent high-resolution coarse-grained model. Our results reveal a strong correlation between condensate viscosity and density, as well as with protein/RNA length and the number of stickers *vs.* spacers in the amino-acid protein sequence. Moreover, we couple the SSRMI and the OS technique to nonequilibrium molecular dynamics simulations that mimic the progressive liquid-to-gel transition of protein condensates due to the accumulation of inter-protein *β*-sheets. We compare the behaviour of three different protein condensates—i.e., those formed by either hnRNPA1, FUS, or TDP-43 proteins—whose liquid-to-gel transitions are associated with the onset of amyotrophic lateral sclerosis and frontotemporal dementia. We find that both SSRMI and OS techniques successfully predict the transition from functional liquid-like behaviour to kinetically arrested states once the network of inter-protein *β*-sheets has percolated through the condensates. Overall, our work provides a comparison of different modelling rheological techniques to assess the viscosity of biomolecular condensates, a critical magnitude that provides information on the behaviour of biomolecules inside condensates.

## I. INTRODUCTION

Biomolecular condensates are membraneless assemblies that contribute to the spatiotemporal organisation of biomolecules in the cytoplasm and the nucleoplasm ^1–6^. These condensates—mainly formed by multivalent proteins and nucleic acids^7,8^— actively participate in numerous aspects of cell function, such as in compartmentalisation ^6,9–13^, genome organisation ^14–17^, gene expression^14,18,19^, formation of super-enhancers^20^, cell signaling^2,21^, buffering of protein concentrations^22^, the sequestration of harmful components in the cell^23^ and many others^24–28^. Biomolecular condensates are thought to form via the process of liquid-liquid phase separation (LLPS), which refers to the physicochemical demixing of a biomolecular mixture into different coexisting liquid phases with distinct concentrations^3^. Microscopically, liquid-like behaviour within phase-separated condensates originates on weak multivalent attractive interactions that proteins and nucleic acids can establish^29^. Such weak and transient intermolecular interactions translate into dynamic binding and unbinding, free molecular diffusion within condesates, and facile exchange of species in and out of the condensates^15,16^. Initially, the liquid-like behaviour of the molecules within the condensates was thought to be a defining feature of such systems. However, more recently, the material properties of biomolecular condensates have been recognised as more diverse than initially anticipated, with condensates encompassing low to high viscosity fluids^30,31^, hydrogels^32,33^, and even solid-like states^34,35^.

Whilst the liquid-like behaviour of the condensates seem to underpin their functions during health^36,37^, kinetically trapped states are often associated to the proliferation of multiple neurodegenerative disorders^38^—such as amyotrophic lateral sclerosis (ALS)^33^, Parkinson’s^39^, Alzheimer’s^40^, or frontotemporal dementia (FTD)—as well as to certain types of cancers^41^ and diabetes^42^. Several factors that have been proposed as key drivers for condensate liquid-to-gel/solid transitions include altered salt-concentration or temperature^30,43^, post-translational modifications^40,44^, protein mutations^45–47^, and most prominently, protein folding and misfolding events^13,48–52^. All these factors are expected to favour progressive condensate rigidification by increasing the binding affinity among species, and therefore, slowing down the timescales of inter-protein unbinding events.

To characterize the progressive rigidification of condensates that initially display liquid-like behaviour and gradually change their material properties into gels or soft glasses (i.e., ‘maturation’), several experimental techniques including fluorescence recovery after photo bleaching (FRAP), green fluorescence protein (GFP) FRAP, fluorescence correlation spectroscopy, or active microrheology have been successfully employed^12,30,53–56^. Further, viscoelastic properties such as viscosity (*η*) have been also measured through passive microrheology techniques (i.e., bead-tracking)^57–63^, in which the trajectory of the beads can be registered and their mean squared displacement (MSD) calculated. Then, the droplet viscosity can be inferred from the diffusion coefficient obtained through the MSD by using the Stokes-Einstein relation^64^. Matured condensates usually exhibit reduced fusion propensities and longer recovery times after photo-bleaching^4,53,54,65–70^, which suggest that the diffusion of molecules within the condensate is significantly reduced. While viscoelastic measurements allow us to identify the gradual transition of functional condensates into pathological aggregates, they are not sufficient on their own to uncover the underlying molecular mechanisms of such transitions. Rationalizing from a microscopic perspective the dysregulation of condensates into pathological aggregates is fundamental to devise effective strategies to prevent condensate age-related diseases^33^ such as neurodegenerative disorders^71^ and some types of cancer^72^.

In that respect, computer simulations are a powerful tool to uncover the molecular mechanisms that explain the changes in viscosity within biomolecular condensates over time^31,51,63,73–76^. From atomistic force fields^77–82^ to coarse-grained (CG) models^83–92^, including lattice-based simulations^93–95^ and mean-field theory^96–98^, computer simulations have significantly contributed to elucidating factors behind condensate phase-separation such as protein and RNA length^99–101^, amino acid patterning^91,102–105^, multivalency^35,106–109^, conformational flexibility^89,110^ or multicomponent composition^90,111,112^. Remarkably, coarse-grained models have uncovered the impact of enhancement of inter-protein interactions in condensate rigidification^51,74^, as well as the formation of kinetically-arrested multiphase condensates from single-component droplets^73,113^. Nevertheless, further insights on the molecular driving forces behind condensate maturation—for instance triggered by inter-protein disordered-to-order structural transitions^50,75,114^, amino acid sequence mutations^115^ or relevant variations on the applied thermodynamic conditions^116^—are urgently needed.

In this work, we apply three different computational methods to evaluate the viscoelastic behaviour of biomolecular condensates formed by proteins and RNA. These methods are the integration of the shear stress relaxation modulus (SSRMI)^117,118^, the oscillatory shear (OS) technique^119,120^, and passive microrheology bead tracking (BT)^30,121–123^. Although these are well known in the field of polymer physics^117,124^, here we test them for the first time in the context of biomolecular condensates and progressive condensate maturation. First, we assess their performance in terms of statistical uncertainty, computational efficiency, and implementation cost using a simple intrinsically disordered protein (IDP) coarse-grained model. Importantly, we find that the three methods provide consistent results for condensate viscosities under different conditions. However, the performance in terms of computational efficiency and statistical error is significantly poorer for the BT technique. Then, we apply the SSRMI and OS techniques for determining the droplet viscosity of a set of 7 different IDPs and 5 peptide/RNA complex coacervates using a sequence-dependent high-resolution coarse-grained model^102,125,126^. Remarkably, in all cases, the agreement between the SSRMI and OS methods to evaluate viscosity is reasonable. Furthermore, we identify a clear correlation between the condensate viscosity and IDP length, as well as with the number of stickers *vs.* spacers^127^ in the amino-acid protein sequence when viscosity is measured at a constant ratio of temperature (T) over the critical temperature of each system 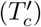. However, when temperature is kept constant instead of 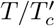, viscosity correlates with condensate density and critical temperature. Lastly, by means of these two techniques, we track the progressive maturation of three of the most relevant protein low-complexity domains related to the onset of ALS and FTD, which are the heterogeneous ribonucleoprotein A1 (hnRNPA1)^13,114,128^, fused in sarcoma (FUS)^49^, and the TAR DNA-binding Protein of 43 kDa (TDP-43)^50,129^. We find that both the shear stress relaxation modulus and oscillatory shear techniques predict the transition from liquid-like behaviour to a gel-like state once the inter-molecular network of *β*-sheets has fully percolated through the condensate. Such percolation of strong *β*-sheets connections frustrates the longtime self-diffusion of proteins within condensates. Taking together, our study provides an evaluation of modelling rheological techniques that enable assessment of different possible scenarios in which biomolecular condensates may, or may not, rigidify over time.

## II. RESULTS

### A. An IDP coarse-grained model for benchmarking viscosity calculations through different techniques

The viscosity is a fundamental time-dependent material property of condensates that emerges from the internal friction of proteins within, and thus, changes from the early stages of condensate nucleation to its maturation over time^30^. Despite its importance, the estimation of the condensate viscosity through computer simulations is not routinely done^126,130^. Here, we test the validity and computational performance of three different numerical methods to compute viscosity. To start, we employ a simple coarse-grained model for phase-separating IDPs^100^. In this model, each IDP consists of a flexible polymer of *N* = 50 beads, where each bead represents a group of several amino acids. We mimic the ability of phase-separating IDPs to establish numerous weak and promiscuous protein-protein interactions at short molecular distances with a short-ranged attractive Lennard-Jones (LJ) potential among non-bonded protein beads:

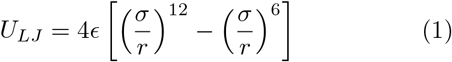

where *σ* accounts for the molecular diameter of each bead, *r* is the inter-bead distance, and *ϵ* defines the maximum attractive interaction among different beads. The LJ potential approximates the various types of protein–protein interactions driving LLPS (e.g., hydrophobic, electrostatic, cation-*π*, and *π*-*π*)^29,89^. For computational efficiency, the pair force computed from the gradient of *U_LJ_* is truncated to zero at a cut-off distance of *r_c_* = 3*σ* so that non-bonded forces act only between pairs of particles with *r* < *r_c_* ^100^. To account for the covalent bonds among subsequent groups of amino acids within a given IDP, consecutive beads are joined together with a stiff harmonic potential, *U*_Bond_, of the following form:

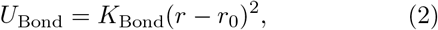

where *K*_Bond_ controls the stiffness of the bond and *r*_0_ is the equilibrium bond length. The model presents a spring constant *K*_Bond_ = 7.5 · 10^4^*ϵ*/*σ*^2^, and equilibrium bond length corresponding to 1*σ*. Non-bonded interactions between adjacent beads directly connected are excluded. Furthermore, for computational efficiency, the solvent is modelled implicitly; hence the protein-poor liquid phase corresponds to a vapour phase and the protein-rich liquid phase (or the condensate) to a liquid phase. For this model, we define the following magnitudes in reduced units: temperature as *T*\* = *k_B_T/ϵ*, number density as *ρ*\* =(N/V)*σ*^3^, pressure as *p*\* = *pσ*^3^/*ϵ*, and reduced time (*τ*) as 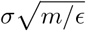; being *ϵ, σ*, and *m* equal to 1. The phase diagram in the *T*\*–*ρ*\* plane for our IDP model obtained through Direct Coexistence (DC) simulations^131^ is presented in Fig. 1(a). Further details of the simulation system sizes and the DC method are provided in the Supplementary Material (SM).

**FIG. 1.**
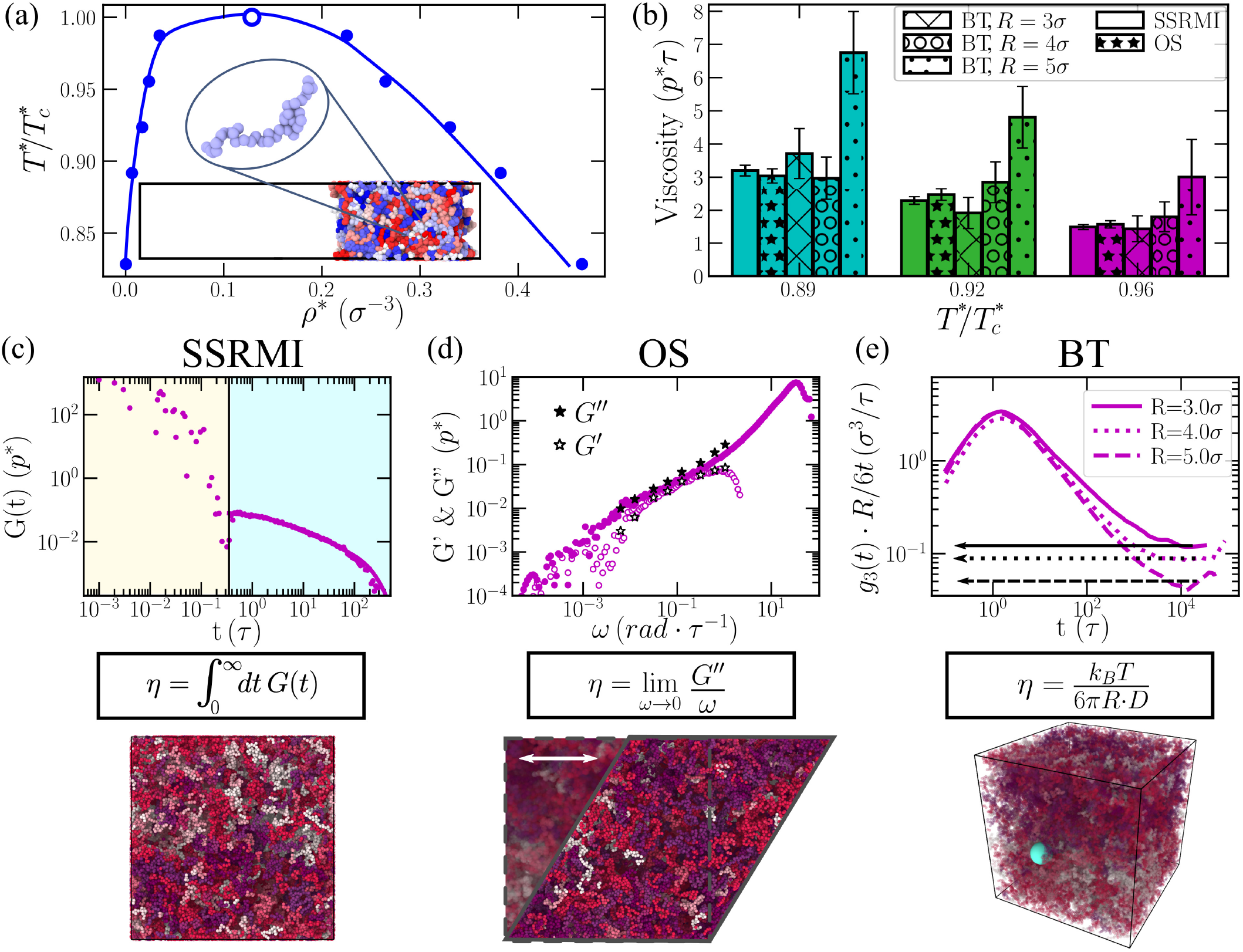
Applied computational methods to evaluate the viscosity in biomolecular condensates. (a) Phase diagram in the *T*\*–*ρ*\* plane for our IDP coarse-grained model using 50-bead chains obtained through Direct Coexistence simulations^131^. Filled circles indicate the coexisting densities obtained from DC simulations (the inset shows a phase-separated condensate in a DC simulation) whereas the empty circle accounts for the system critical temperature 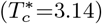 obtained through the law of rectilinear diameters and critical exponents^132^. (b) Condensate viscosity at different temperatures obtained through SSRMI, OS, and BT calculations as indicated in the legend. For the BT technique, we include results with different probe bead radii as specified in the legend. (c) Top: Shear stress relaxation modulus as a function of time for an IDP condensate at 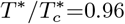. The vertical black line separates the timescale corresponding to the computed term via numerical integration at short-times and the part evaluated via the Maxwell-modes fit at long timescales. Middle: General equation to obtain viscosity through the SSRMI method. Bottom: IDP condensate simulation box in the canonical ensemble (at the condensate coexisting density) employed to compute *G*(*t*). Different IDPs are coloured with different tones (as in (d) and (e) Bottom Panels). (d) Top: Elastic (*G′*) and viscous (*G″*) moduli as a function of frequency (*ω*) from OS calculations (filled and empty stars respectively) and from SSRMI simulations (filled and empty purple circles respectively) at 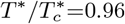. Middle: General equation to obtain viscosity through the OS technique. Bottom: IDP condensate simulation box in the canonical ensemble (at the condensate coexisting density) after applying a shear deformation (*γ_x,y_*). (e) Top: Mean-squared displacement (referred as *g*3(*t*)) of an inserted bead within an IDP condensate 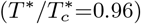 multiplied by *R*/6*t* (referring *R* to the bead radius and *t* to time) as a function of time for beads with different radii as indicated in the legend. The plateau at long timescales (denoted by horizontal lines) shows the value of *R* · *D* (being *D* the diffusion coefficient) at the diffusive regime. Middle: Stokes-Einstein equation for computing viscosity through the BT method. Bottom: IDP condensate simulation box in the canonical ensemble (at the condensate coexisting density) containing a single-bead with a radius of 5*σ* (green sphere).

#### 1. Shear stress relaxation modulus integration (SSRMI)

To begin, we evaluate the viscosity of a condensate of our IDP model by means of the SSRMI method. The time-dependent mechanical response of a viscoelastic material when it is subjected to a small shear deformation can be described by the shear stress relaxation modulus (*G*(*t*))^117^. The shear viscosity (*η*) can be straightforwardly calculated by integrating the shear stress relaxation modulus in time:

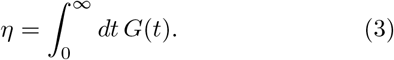

In the limit of zero deformation, *G*(*t*) can be determined by computing the auto-correlation of any of the off-diagonal components of the pressure tensor at equilibrium^124,133–135^:

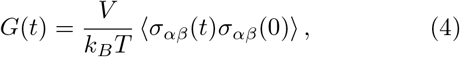

where *σ_αβ_* is an off-diagonal component (*αβ*) of the stress tensor, *V* is the volume, and the correlation average is taken at equilibrium over all possible time origins. Nevertheless, if the system is isotropic, a more accurate expression of *G*(*t*) can be obtained by using the six independent components of the pressure tensor, as shown in Refs.^118,124^:

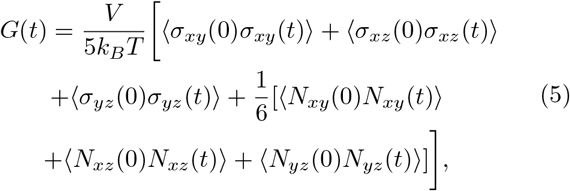

where *N_αβ_* = *σ_αα_* – *σ_ββ_* is the first normal stress difference. This correlation can be easily computed on the fly during a simulation, with no significant CPU cost and no need to post-process the trajectory. For instance, in the LAMMPS Molecular Dynamics (MD) package, this can be done by using the compute ave/correlate/long in the USER-MISC package^136^. To avoid the typical noisy nature of the relaxation modulus in the terminal decay region obtained in protein condensate simulations^75,130^, we follow a particular strategy to estimate the viscosity. Whilst at short timescales *G*(*t*) is smooth and the integral can be computed using numerical integration (Eq. 3), at longer timescales *G*(*t*) is fitted to a series of Maxwell modes (*G_i_* exp(–*t/τ_i_*)) equidistant in logarithmic time^117,137^ and then, the function is integrated analytically. Therefore, viscosity is effectively obtained by adding two different terms

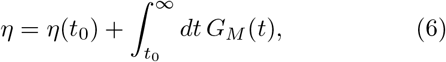

where *η*(*t*_0_) corresponds to the computed term for short time-scales, *G_M_*(*t*) = ∑^*M*^ *G_i_* exp(–*t/τ_i_*) is the part evaluated via the Maxwell modes fit at long time-scales, and *t*_0_ is the time that separates both (i.e., black vertical line in Fig. 1(c)).

The SSRMI method is exact, within the accuracy of the underlying simulation, and gives the right value of *G*(*t*) in the limit of zero deformation (*γ* → 0). A similar measurement can be performed experimentally by applying a shear deformation *γ* and measuring the evolution of the stress response *σ_xy_*(*t*) to determine the stress modulus as *G*(*t*) = *σ_xy_*(*t*)/*γ*. Hence, the relaxation modulus *G*(*t, γ*) can be experimentaly estimated, despite the signal of *σ_xy_*(*t*) being weaker in the small amplitude regime, which is necessary to maintain the linear viscoelastic region. The SSRMI method has been recently applied by us^75,130^ to evaluate viscosities in phase-separated condensates of RNA-binding proteins both in absence and presence of RNA. One of the main advantages of the direct evaluation of *G*(*t*) from simulations is that it provides critical information not only on how the material properties of condensates may change upon maturation, but also on how such changes are dictated by different relaxation mechanisms of the proteins that compose them (Fig. 1(c)). While at short timescales (beige region; Fig. 1(c)), the stress relaxation modulus is mostly dependent on the formation and breakage of short-range interactions and on intramolecular reorganisation (i.e., intra-molecular protein conformational fluctuations, such as bond or angle relaxation modes), at long timescales (light blue region; Fig. 1(c)), the stress relaxation modulus is mainly dominated by inter-molecular interations, long-range conformational changes (i.e. protein folding/unfolding events), and protein diffusion within the crowded liquid-like environment of the condensate. Moreover, the calculation of *η* through this method does not depend on the size of the system, apart from the obvious limit to avoid finite size effects. As the system grows in size, the equilibrium value of the shear stress goes to zero, and the fluctuations become smaller. However, the size of the fluctuations of *σ_xy_* decay with 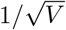, and therefore the calculation of *G*(*t*) becomes independent of *V*.

In Fig. 1(c), we show the time evolution of *G*(*t*) (purple circles) measured for the IDP model condensate at the coexisting density corresponding to 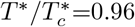. By numerical integration (beige region) and analytical integration (light blue region) *G*(*t*), as shown in Eq. 6, we can obtain the condensate viscosity, as depicted in Fig. 1(b) for different temperatures.

#### 2. Oscillatory shear (OS) technique

The second approach that we employ to determine the viscosity of phase-separated condensates is the OS method. In this approach, a sinusoidal strain with angular frequency *ω* is applied to the condensate in simple shear:

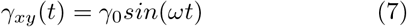

where *γ_xy_*(*t*) = Δ*L_x_/L_y_*(*t*) represents the shear deformation applied to the simulation box in the x direction relative to the box dimension *L_y_*, and *γ*_0_ is the amplitude of the imposed deformation (Fig. 1(d)). Please note that the box is cubic so that *L_x_* = *L_y_* = *L_z_* ≡ *L*. This type of deformation can be easily applied to the simulation box by using the ‘fix deform’ command of the LAMMPS package, with the option ‘wiggle’. If the biomolecular condensate lies within the linear viscoelastic (LVE) regime, then the stress response will be

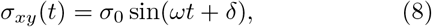

where *σ*_0_ refers to the amplitude of the response and *δ* to the phase shift angle. Within the LVE regime, the ratio *σ*_0_/*γ*_0_ is constant and the shear response presents a sinusoidal shape. To determine the optimal *γ*_0_, an amplitude sweep is needed to ensure that the shear deformation is within the LVE regime, but also that the stress response signal with such deformation is detectable (see SM for further technical details). Once selected an amplitude within the LVE regime (*γ*_0_; which in our simulations is usually 0.6*L*, where *L* is the size of the initial cubic box), we perform a frequency sweep (avoiding high frequencies to prevent overheating; please note that the maximum frequency should be sufficiently smaller than the inverse of the characteristic relaxation time of the thermostat), and we measure the transient stress tensor response in the direction of the oscillatory shear (*σ_xy_*(*t*)). Then, by fitting the stress response for all frequencies (after 20 periods of sampling), we can calculate *σ*_0_ and *δ*, and thus the elastic modulus (*G′*, also known as storage modulus) and viscous modulus (*G″*, also known as loss modulus) through the following expressions:

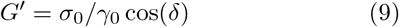

and

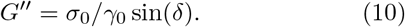

By means of the OS technique, the viscosity can be estimated in the limit of^117^

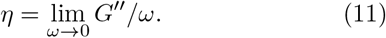

For computing the viscosity of a liquid in this regime (where *G″* ∝ *ω* and *G′* ∝ *ω*^2^)^117^, long simulations using large amplitudes are required so that the stress response is higher than the fluctuations of the system. Furthermore, from the representation of *G′* and *G″* as a function of *ω*, the viscoelastic behaviour of the system can be inferred. If *G′* > *G″*, elasticity dominates over flow, and hence the system exhibits solid-like behaviour. On the contrary, the viscoelastic response of a liquid is markedly different. The terminal response of a liquid condensate is dominated by the loss modulus because the stress is nearly in phase with the shear rate (the time derivative of the applied shear deformation *γ_xy_*(*t*)), and hence *G″* is higher than *G′* at intermediate frequencies. Despite the fact that this technique has been experimentally applied to numerous soft matter and polymeric systems^119,120^ (some of them including even chocolate^138^ or mozzarella^139^), its application to protein condensates has been much more limited^30,140^, mainly due to sample size requirements (bulk rheology measurements need, at least, volumes of the order of milliliters). In computer simulations, the OS technique has been mainly employed to characterise polymeric systems^117,123,141–143^.

In Fig. 1(d), we show the values for *G′* (empty stars) and *G″* (filled stars) as a function of the applied frequency for a condensate of our IDP coarse-grained model using the OS technique. As can be seen, excellent agreement is obtained with the results from the SSRMI method for *G′* (purple empty circles) and *G″* (purple filled circles). Furthermore, when viscosity is estimated through OS by means of Eq. 11, a good agreement is also found with SSRMI predictions for IDP condensates at different temperatures (Fig. 1(b)).

#### 3. Bead tracking (BT) method

The bead tracking method is a passive microrheology technique widely used in experiments to determine the viscosity of a given material^117,144^. For biomolecular condensates, this is the technique that has been mainly used to measure the viscosity of *in vitro* phase-separated droplets displaying both liquid-like^59,61,62,140,145^ and gel-like behaviour^30,121,122^. The idea behind this method is as simple as introducing passive probe spherical beads (with a typical radius, R, of the order of hundreds of nanometres^144^), and measure their mean square displacement (MSD(t) = 〈(*r*(*t*)–*r*(0))^2^)〉 from which the diffusion coefficient (*D*) of the bead can be calculated as^146,147^:

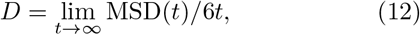

where the limit indicates the time when the diffusive regime is attained. Importantly, the bead size needs to be larger than the characteristic mesh size of the system. Otherwise, the probe would move freely without experiencing the force of network strands or entanglements^117^. Then, using the Stokes-Einstein relation^148^, the viscosity of the medium can be calculated:

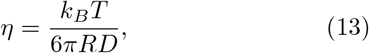

where *k_B_* refers to the Boltzmann constant and *T* to the system temperature. The BT technique is highly suitable for characterising the viscoelastic properties of biological systems, such as biomolecular condensates^30,59,61,62,121,122,145^, as it can be performed in volumes of the order of *μ*l. Importantly, microrheology BT can also be performed *in vivo* by tracking the motion of micrometre-sized beads (or even smaller beads^149^) inserted inside cells, as performed in the cytoplasm of developing C. *elegans* embryos^150^. Despite microrheology bead tracking can be active (i.e., when the particle is moved in the medium by means of optical tweezers or magnetic forces^30^), here we shall focus on passive BT where only thermal energy drives the probe particles across the medium exerting minimal deformation, and the motion of the particle can be related to the mechanical properties of the medium.

In our simulations, we perform passive single-particle bead tracking to calculate the viscosity of the condensates via the Stokes-Einstein relation (Eq. 13). The probe particles are modelled with an Ashbaugh-Hatch potential^151^ of the following form:

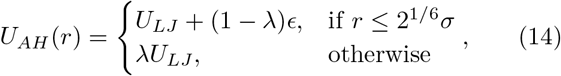

where *U_LJ_* refers to the standard LJ potential presented in Eq. 1, *ϵ* to the LJ potential depth (set to *ϵ*=4 for the probe bead to ensure no-slip boundary conditions), and *λ* is a scaling factor that modulates the degree of attraction between probe beads and IDPs (where *λ* = 0 establishes a purely repulsive interaction and *λ* = 1 a standard LJ interaction). The mass of the probe bead is set to *m* =1 and the cut-off distance for the *U_AH_* interaction is 3 times the probe bead molecular diameter. We explore bead particles with radii of 3, 4, and 5*σ* (referring *σ* to the molecular diameter of the residue beads in our IDP coarse-grained model). For both bead–bead self-interactions and bead–IDP cross-interactions, we set a value of *λ* equal to zero. In this limit, the Ashbaugh-Hatch potential is identical to a Weeks-Chanled-Andersen^152^ potential, and hence the inserted probe beads act as pseudo-hard-sphere particles with the surrounding media^153^. Please note that despite self-interactions among probe beads are set to be purely repulsive, we only study this technique using a single probe bead (see the SM for further technical details).

Following previous work on bead tracking simulations for polymeric systems^123^, we introduce a probe bead (with radius of 3, 4, and 5*σ*) within phase-separated condensates under bulk conditions (Fig. 1(e) Bottom panel). By plotting the mean square displacement (*g*_3_(*t*) ≡ MSD(*t*)) multiplied by *R*/6*t* as a function of time, we can identify the timescale at which the diffusive regime is attained (i.e. the function reaches a plateau) and introduce the value of such plateau (*R · D*) in the Stokes-Einstein equation to obtain *η* (Eq. 13). In Fig. 1(e; Top panel), we depict by black horizontal lines the value of *R · D* on the plateau for different bead radii at 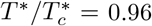. Although the values of *R · D* slightly depend on the bead radius, the BT method predicts the same viscosity within the uncertainty as the SSRMI and OS methods at 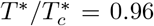 (Fig. 1(b)). Nevertheless, such agreement between BT, OS, and SSRMI methods to predict *η* is only observed at low temperatures when probe beads of *R* = 3*σ* and 4*σ* are introduced within the condensates (Fig. 1(b)). The reason behind that fact is the much longer timescales required to reach the diffusive regime at low temperatures with very large beads (i.e., *R* = 5*σ* or 6*σ*; see Fig. S1), which entails a huge computational effort to observe a smooth plateau from which a reliable value of *η* can be obtained (as it easily occurs for all bead sizes at high temperatures). Moreover, since the probe bead size must be significantly larger than the characteristic mesh size of the system^117,123^, lowering the bead size below 3*σ* to increase its diffusion would lead to an underestimate of the condensate viscosity (as shown in Fig. S1). Hence, we note that the BT method can only be safely applied in low viscous condensates (or at high temperatures) where the large sampling of the bead trajectory guarantees consistent results independently of the inserted bead size (Fig. 1(e); purple bars).

Therefore, despite being conceptually a straightforward approach, the bead tracking method requires long simulation timescales to ensure that the mean square displacement of the inserted beads are properly sampled. The computational efficiency of the bead tracking method, and thus its associated statistical error, is significantly hampered by the requirements to include only one bead in the simulation box (or at least the concentration of beads must be low enough to guarantee that beads do not interact with each other) and to use probe beads that are larger than the characteristic mesh size. However, it can still provide reasonable estimations of *η* as shown in Fig. 1(b) under certain conditions (i.e., within low viscous media).

### B. Condensate viscosity is fundamentally determined by protein/RNA length, stickers abundance, and condensate density

Once the different advantages and drawbacks of the SSRMI, OS, and BT methods have been discussed in determining the viscosity of the condensate, we move away from generic proteins and now explore the dependence of the viscosity of 12 different phase-separated protein/RNA condensates using a sequence-dependent coarse-grained model. We compare the changes in viscosity among these systems by focusing on the protein length, amino acid sticker abundance, condensate density, molecular system mass, critical temperature, and number of charged residues along the sequence. Specifically, we use the reparameterization^125^ of the residue-resolution HPS model from the Mittal’s group^103^. We have recently shown that the HPS-Cation-*π* reparameterization qualitatively reproduces the relative propensity of numerous RNA-binding proteins to phase separate under physiological conditions^130^, as well as their RNA-concentration-dependent re-entrant phase behaviour^154–157^. Within this force field, hydrophobic and cation–*π* interactions are modelled through shortrange pairwise potentials, and electrostatic interactions through a Yukawa/Debye-Hückel long-range potential. Additionally, bonded interactions between subsequent amino acids (or nucleotides) of the same protein (or RNA) are restrained by a harmonic potential. Moreover, within the HPS-Cation-*π* force field the solvent is implicitly modelled by the screening length of the Yukawa/Debye-Hückel potential, which is tuned to reproduce protein phase behaviour at physiological salt concentration (~150 mM). All details regarding the force field parameters and simulation setups are provided in the Supplementary Material.

The set of phase-separating biomolecules that we explore includes the following IDPs: the DEAD-box helicase 4 (Ddx4)^158^, *α*-synuclein^159^, the microtubule-associated neuronal IDP Tau K18^160^, the arginine-glycine rich-region of LAF-1 (LAF-1-RGG)^81^, and the low complex domains (LCD) of the heterogeneous nuclear ribonucleoprotein A1 (A-LCD-hnRNPA1)^13^, fused in sarcoma (FUS-LCD)^30^, and the TAR DNA-binding Protein of 43 kDa (TDP-43-LCD)^129^. Moreover, we investigate the viscoelastic behaviour of 5 charge-matched complex coacervates (which can only phase-separate via heterotypic interactions^101^): the proline-arginine 25-repeat dipeptide (PR25) in presence of single-stranded disordered polyUridine RNA (polyU)^80^ of 50-nucleotides (polyU50) and 100-nucleotides length (polyU100); and three 50:50 binary mixtures of polyU and poly-Arginine (polyR)^60^ with different chain lengths, polyR50/polyU50, polyR50/polyU100, and polyR100/polyU100.

We first evaluate the phase diagram (through DC simulations^131^; see SM for further details) of the entire set of intrinsically disordered proteins (Fig. 2(a)) and complex coacervates (Fig. 2(b)). Remarkably, we find that the HPS-Cation-*π* coarse-grained model qualitatively predicts the higher ability to phase separate for IDPs such as Ddx4^162^ or A-LCD-hnRNPA1^53,163^ compared to other low-complexity domains such as FUS-LCD^29,155^ or TDP-43-LCD^164^, which require a higher protein saturation concentration in experiments. In contrast, the high critical saturation concentration of *α*-synuclein to undergo LLPS^159^, which is similar to that of FUS-LCD^29,155^ is not qualitatively well predicted by the model. For the complex coacervates, we observe that for the same peptide/RNA length (i.e., PR_25_/polyU50 vs. polyR50/polyU50, and PR_25_/polyU100 vs. polyR50/polyU100), polyR/polyU condensates always display higher ability to phase separate, in agreement with experimental *in vitro* findings^60,80^. Furthermore, as expected, we also reproduce the higher ability to undergo LLPS as the length of the peptides and RNA increases in both families of complex coacervates (Fig. 2(b)).

**FIG. 2.**
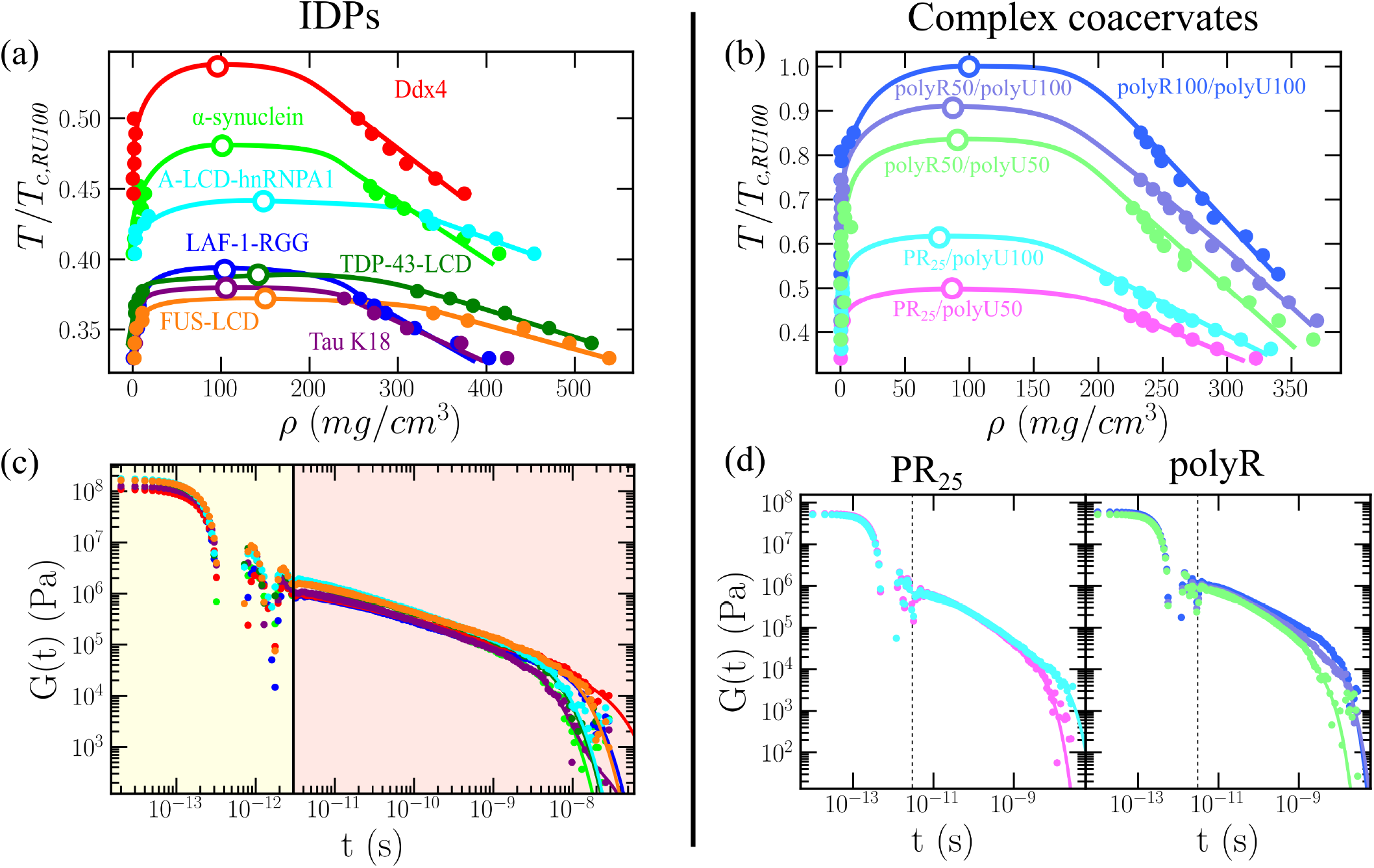
Phase diagram and shear stress relaxation modulus for a set of IDP/polyU phase-separated condensates. (a) Phase diagram in the T–*ρ* plane for Ddx4, *α*-synuclein, Tau K18, LAF-1-RGG, A-LCD-hnRNPA1, FUS-LCD, TDP-43-LCD using the HPS-Cation-*π* force field^103,125^. (b) Phase diagram in the T–ρ plane for PR25/polyU50, PR25/polyU100, polyR50/polyU50, polyR50/polyU100, and polyR100/polyU100 using the HPS-Cation-*π* force field^103,125^. In both panels (a) and (b), filled symbols represent the coexistence densities obtained via Direct Coexistence simulations^131^, while empty symbols depict the estimated critical points by means of the law of rectilinear diameters and critical exponents^161^. Moreover, temperature has been renormalized by the critical temperature (*T*_c,RU100_ ≡ *T*_c,polyR100/polyU100_) of the system with highest *T_c_*, which is the charge-matched polyR100/polyU100. The statistical error is of the same order of the symbol size. (c) and (d): Shear stress relaxation modulus *G*(*t*) of the systems shown in panel (a) and (b) at 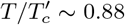 (referring 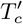 to the critical temperature of each system) and at the bulk condensate density at such temperature. The vertical continuous (c) and dashed (d) lines separate the timescale corresponding to the computed term via numerical integration at short timescales and the part evaluated via the Maxwell-modes fit at long timescales (Eq. 6).

Next, we evaluate the condensate viscosity for all systems employing the SSRMI (Figs. 2(c) and 2(d)) and OS (Fig. 3(a)) methods. The conditions at which we undertake these calculations are at 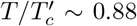 (referring 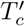 to the critical temperature of each system) and the condensate bulk density at such temperature. Since in Fig. 1(b) we show that the BT method only provides consistent estimations of *η*, independently of the probe bead size at relatively high temperatures or unless extremely long simulations are performed, for this set of biomolecular condensates we only carried out SSRMI and OS calculations. In Figs. 2(c) and 2(d) we show the shear stress relaxation modulus as a function of time for all IDPs and complex coacervates, respectively. In our simulations, all of the condensates are able to relax (i.e., the correlation function decays) at long timescales and therefore exhibit liquid-like behaviour. Moreover, in Fig. 3(a), we report the elastic (G’; empty stars) and viscous (G’’; filled stars) moduli as a function of frequency from our OS calculations. The agreement in all cases for the entire regime of frequencies studied between OS and SSRMI simulations is exceptional. The results of *G′* and *G″* from SSRMI calculations have been obtained by applying the Fourier transform using the open source RepTate software^165^. In accordance with the *G*(*t*) decays observed in Figs. 2(c) and 2(d), the values of G’’ are higher than G’ within the moderate frequency regime for all systems, further supporting that the viscoelastic behaviour of the condensate is dominated by the loss modulus (e.g., liquid-like behaviour) given that the stress is nearly in phase with the shear rate.

**FIG. 3.**
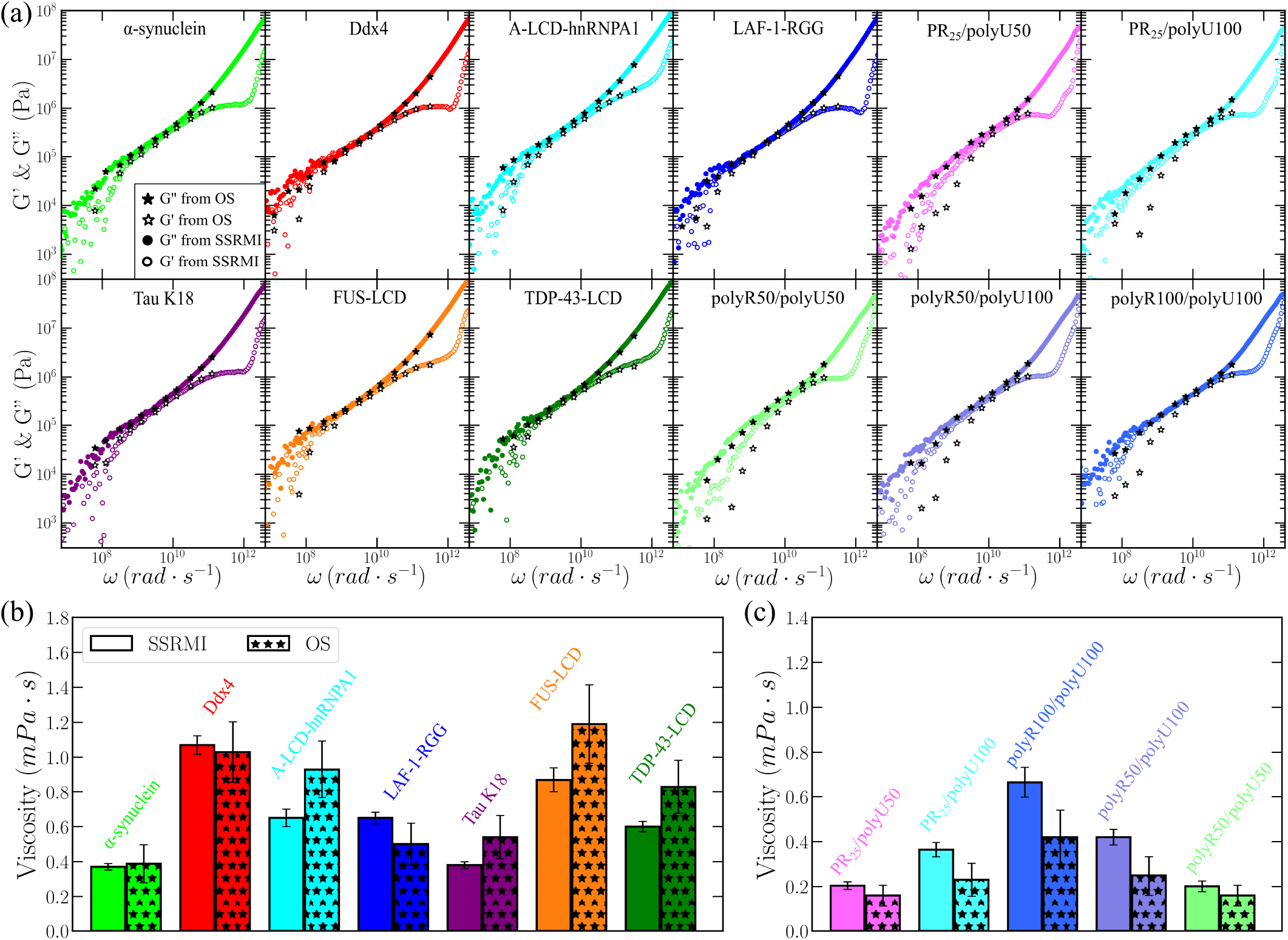
Condensate viscosity for different IDP and RNA complex coacervates evaluated through the integration of the shear stress relaxation modulus (SSRMI) and the oscillatory shear (OS) method. (a) Elastic (*G′*) and viscous (*G″*) moduli as a function of frequency (*ω*) from OS calculations at 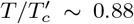 (filled and empty stars respectively) and from SSRMI simulations (filled and empty coloured circles respectively) for different IDP condensates and complex coacervates as indicated in the legend.(b) Viscosity computed via the SSRMI and OS methods for the distinct IDP condensates. (c) Viscosity obtained through the same two methods for the different complex coacervates.

By integrating in time the shear stress relaxation modulus (Eq. 6) shown in Figs. 2(c) and 2(d) for the distinct IDPs and complex coacervates, we can obtain the viscosity of the condensate. Moreover, taking the limit of Eq. 11 to very low frequencies, we can also evaluate *η* using the OS approach. In Figs. 3(b) and 3(c), we report the viscosities obtained through the two methods for the set of IDPs and complex coacervates studied, respectively. It can be noted a fair agreement in the predicted viscosity between the OS and SSRMI methods for most of the studied biomolecular condensates. Given that *η* is a magnitude that can dramatically vary (orders of magnitude) with small changes in protein intermolecular binding^30,60,74^ or in applied thermodynamic conditions^116,166^, the fact that, for most condensates, the predicted viscosity differs by less than a factor of 1/2, clearly indicates the robustness of our calculations. However, we acknowledge that the viscosity values from the SSRMI calculations are likely more accurate than those using the OS method, since the former do not rely on the limit of *G″/ω* at very low frequencies, where the signal in the stress response is particularly low^117^.

Interestingly, from the results shown in Figs. 3(b) and 3(c) at 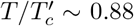, we find a clear correlation between viscosity and protein/RNA length, defined as the number of amino acids or nucleotides in the macromolecule (Fig. 4(a)). As the length of the IDP or RNA chain increases, the viscosity of the phase-separated condensates also augments. Moreover, the same correlation holds for the protein/RNA molecular mass (Fig. 4(b)) since the increase in length directly impacts the molecular weight. However, when plotting *η* against condensate density at 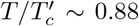 for all systems, we do not observe a clear trend (Fig. S3(e)). That might be a striking result given that a larger density or packing fraction should lead to higher viscosity values^130,166^. Nevertheless, we identify cases such as Ddx4 in which despite presenting a low condensate density, its viscosity is the highest (in correspondence with its length). On the other hand, A-LCD-hnRNPA1 has a moderate length (i.e., 135 residues), its condensates are one the most dense of the set, and its viscosity is just moderate (Fig. S3(e)). We also interrogate the correlation between *η* measured at 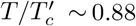 and the relative critical temperature of each system (i.e., ability to undergo LLPS in our model, which is directly inverse to the protein saturation concentration^162^). As shown in Fig. S3(d), there is no clear evidence, according to our simulations, that systems with higher critical temperature should present higher viscosity as long as the conditions at which *η* is determined are equidistant to 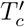 for each system. Nonetheless, when our measurements are carried out at constant *T* = 340K for all the systems instead of 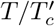 (Fig. 4(e)), a clear dependence is observed between condensate viscosity and critical temperature for the studied set of IDPs and complex coacervates. We hypothesize that this correlation might be also strongly dependent on the type of interactions promoting LLPS, being mostly of electrostatic nature for the complex coacervates, and a combination of hydrophobic, cation-*π*, *π* – *π*, and electrostatic interactions for the IDP set. Furthermore, we find that for a constant temperature (i.e., *T* = 340K) a strong correlation between condensate viscosity and density arises (Fig. 4(f)) in contrast with the results shown at constant 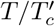 for all systems (Fig. S3(e)).

**FIG. 4.**
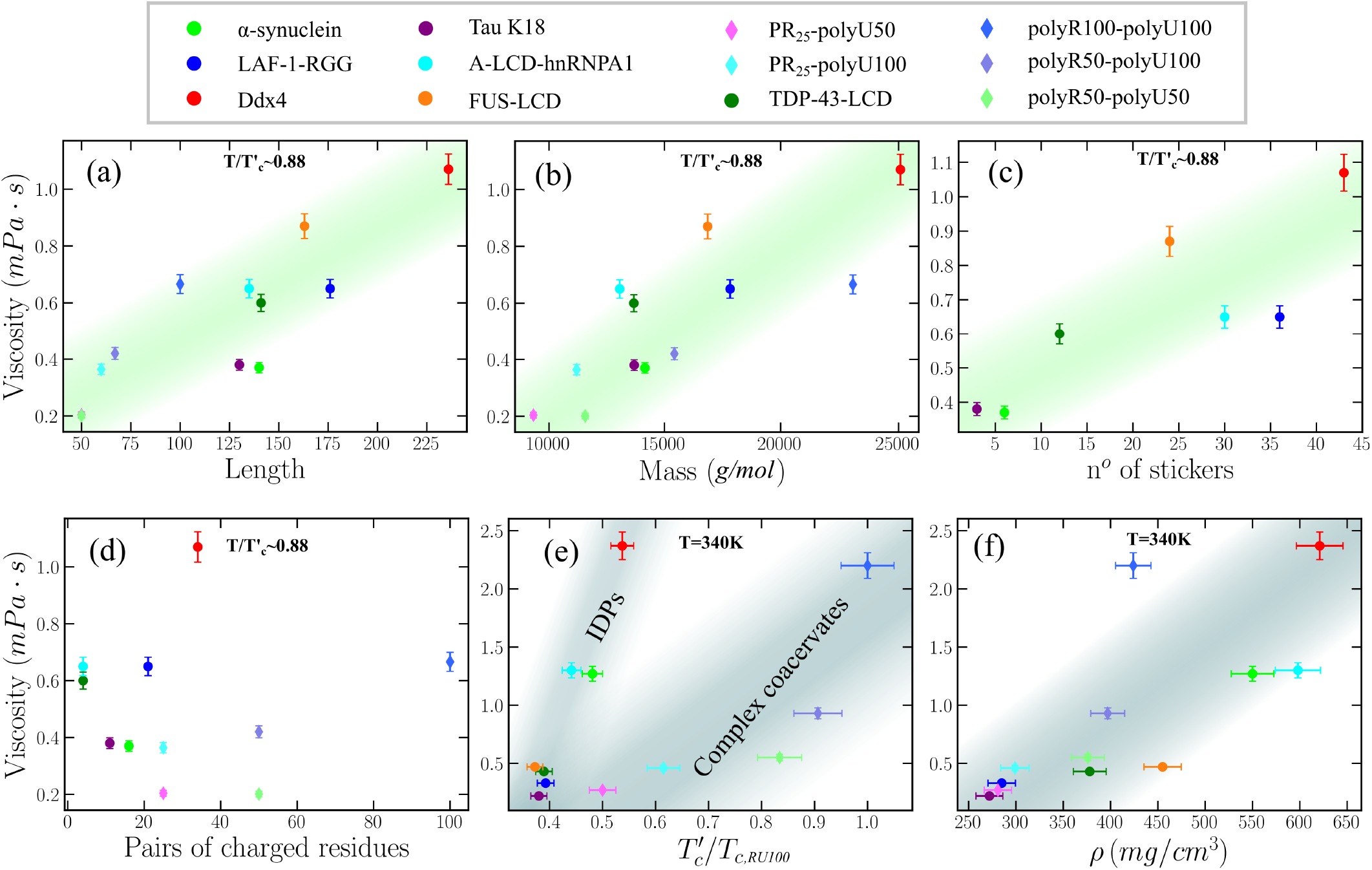
Correlation between condensate viscosity (obtained via the SSRMI method) and chain length (a), molecular weight (b), number of stickers (Y, F and R) across the sequence (c), and number of charged residue pairs of opposite charge (d) at constant 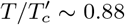 0.88. In contrast, the correlation between viscosity and condensate critical temperature (e), and density (f) is plotted for a constant *T* = 340K. In complex coacervates, we take the average chain length and molecular weight of the two cognate molecules. In panel (e), we note that each critical temperature has been renormalized by the highest critical temperature of the studied set (*T*_c,RU100_, which corresponds to that of polyR100/polyU100). Since the complex coacervates by construction do not contain aromatic residues, their results have not been considered for the correlation shown in panel (c).

We also analyse the correlation of viscosity with the sequence composition across the different studied IDPs (Figs. 4(c) and 4(d)). First, we focus on how *η* is related to the number of stickers and spacers along the different sequences. The framework of stickers and spacers for protein phase separation represents multivalent proteins as heteropolymers made of stickers (i.e. binding sites for associative interactions) and spacers (regions in between stickers)^93,167,168^. Following Ref.^127^, we have considered tyrosine (Y), phenylalanine (F) as main stickers, arginine (R) as a context-dependent sticker, and the rest of the amino acids are classified as spacers. In Fig. 4(c), we show how the viscosity is proportional to the sticker abundance in the different IDPs studied at constant 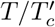. This is an expected result since stickers, due to their stronger intermolecular binding, act as amino acids with an effectively higher friction coefficient, which slows the conformational relaxation of the biomolecules and, thus, increases the viscosity of the condensate. The trend only partially deviates for the case of LAF-1-RGG due to the extremely high abundance of R, which is a context-dependent sticker since, in absence of aromatic residues Y and F, R self-repulsion dominates^127^. Importantly, if we only consider the aromatic residues as LLPS stickers, the correlation between sticker abundance and viscosity becomes poorer than when we also include R as sticker (Fig. S5), highlighting the role of R as a sticker when there is high abundance of aromatics (Fig. 4(c)). When condensate viscosity is plotted against the protein sticker abundance for a constant *T* instead of 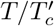, a poorer correlation is found (Fig. S4(c)). We hypothesize that the reason behind this behaviour might be that at constant T, the significant different densities between distinct phase-separating condensates (Fig. 4(f))—in turn regulated by a complex amalgam of sequence features, including the sticker abundance—is mostly controlling the condensate viscosity. Since our simulations are effectively describing the protein phase behaviour at physiological salt concentration, where electrostatic interactions are known to play a key role in sustaining LLPS^169^, we now ask whether a correlation can be found between droplet viscosity and the abundance of charged residues of distinct sign (i.e. total number of pair residues with opposite charge). We show that, very mildly, viscosity might be proportional to the number of charged residues of opposite sign along the studied IDP sequences (at constant 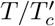 (Fig. 4(d)) and at constant *T* (Fig. S4(f))). However, in both A-LCD-hnRNPA1 and TDP-43-LCD, the number of pair residues with opposite charge is significantly low, and their condensate viscosity is still moderate, showing a much better correlation with the IDP length/molecular weight or the number of stickers throughout the sequence than with the charged residues.

It is important to note that our viscosity results presented in Figs. 3 and 4 generally underestimate the experimental values of *η* for *in vitro* phase-separating condensates^140,170^. The coarse-grained nature of the HPS-Cation-*π* force field in which amino acids and nucleotides are represented by spherical beads in combination with an implicit solvent model, importantly speeds up the system dynamics and leads to an underestimation of approximately 2-3 orders of magnitude in *η*. However, the computational efficiency of the model also enables this type of calculations for phase-separated condensates formed by hundreds of protein replicas^130^. Importantly, with our calculations we recover the experimental observation that increasing polyR and polyU length significantly enhances condensate viscosity, as well as the key role of arginine-uridine interactions in triggering LLPS and increasing viscosity^60^. As shown in Fig. 3(c), arginine-uridine interactions are much more relevant than those of proline-proline, proline-arginine or proline-uridine in regulating condensate viscosity and stability. Furthermore, we recapitulate the observation that within the experimental uncertainty, the viscosity of LAF-1-RGG^170^, TDP-43-LCD^171^, and FUS-LCD^116^ condensates (before maturation) is approximately of the same order. Additionally, it has been reported that Ddx4 inside phase-separated condensates possess an extremely low translational diffusion^158^, our results from Fig. 4 also qualitatively suggests this behaviour. For Tau K18 and *α*-synuclein condensates, to the best of our knowledge, there are no available results for *η*. However, our results for Tau K18 support that phase separation can be experimentally observed only in the presence of molecular crowders, or through complex coacervation with RNA^172^ due to its low abundance of aromatic residues (as shown in Fig. 4(c), purple circle). Hence, these experimental observations^172^ may justify the low viscosity (Fig. 3(b)) and critical temperature (Fig. 2(a)) obtained for Tau K18 condensates.

In summary, although the viscosities predicted by sequence-dependent protein/RNA coarse-grained models^102,125,126,173^ cannot quantitatively match experimental results^140^, they can provide valuable qualitative trends on how the viscoelasitc properties of a given condensate may change upon variations on the thermodynamic conditions^116^, post-translational modifications^174^, mutations^50^, or addition of different cognate molecules^60^. Therefore, establishing robust methodologies to evaluate viscosity via computer simulations can be of great relevance to envision possible strategies to regulate such critical magnitude in the condensate function.

### C. Maturation of protein condensates can be unequivocally tracked by the OS and SSRMI techniques

In this section, we investigate the progressive rigidification of phase-separated condensates due to the gradual accumulation of inter-protein structural transitions over time^13,114,128^. It has been recently shown, both experimentally and computationally, that the interaction landscape of proteins can be significantly transformed by structural transitions^50,73,75,76,114^. The low complexity domains (LCD) of many naturally occurring phase-separating proteins—including FUS^49^, TDP-4^350,129^, or hnRNPA1^13,128^ among many others^114,175^– contain short regions termed Low-complexity Aromatic-Rich Kinked Segments (LARKS), which are prone to forming inter-protein *β*-sheets in environments of high protein concentration^13,116,176^. Although these proteins can form liquid-like condensates, depending on the conditions (i.e., temperature^116^ or concentration^114^), they can also transition into hydrogels over time^47,177,178^. Hence, inter-protein structural transitions have been proposed to trigger transient solidification of, otherwise, liquidlike condensates^45,179^. Importantly, misregulation of biomolecular condensates into solid-like aggregates is associated with the onset of several neurodegenerative diseases^36,180^. Therefore, motivated by these observations, here we explore by means of the SSRMI and OS techniques the impact of transient accumulation of *β*-sheet fibrils in the viscoelastic properties of phase-separated condensates formed by LARKS-containing LCDs of FUS, TDP-43, and hnRNPA1.

For these simulations, we employ an ageing dynamical algorithm recently developed by us^73,75^ to describe the non-equilibrium process of condensate maturation due to inter-peptide *β*-sheet formation. Coupled to the HPS-Cation-*π* residue-resolution model^103,112,125^, our dynamical algorithm approximates the condensate maturation process by considering the atomistic implications (i.e., non-conservative strengthening of inter-protein binding and local protein rigidification^74^) of the gradual and irreversible accumulation of inter-protein *β*-sheet structures in a time-dependent manner, and as a function of the local protein density within phase-separated condensates. In practise, our dynamical algorithm triggers transitions from disordered LARKS to inter-protein structured *β*-sheets when the central *C_α_* bead of a LARKS is in close contact (within a cut-off distance of ~8Å) with three other LARKS of neighbouring proteins^49,50,114^. Therefore, every 100 simulation timesteps, our algorithm evaluates whether the conditions around each fully disordered LARKS are favourable to undergo an ‘effective’ disorder-to-order *β*-sheet transition. In our model, the structural transition is recapitulated by enhancing the interaction strength of four LARKS-LARKS peptides based on the results of our atomistic potential-of-mean force simulations^73,75,76^. In the atomistic simulations, we estimate the binding free energy difference between disordered interacting LARKS peptides, and interacting LARKS peptides that are forming interprotein cross *β*-sheets. We estimate these changes for the three FUS LARKS within the LCD (37SYSGYS42, _54_SYSSYGQS_61_, and _77_STGGYG_82_^49^,^73^), the A-LCD-hnRNPA1 LARKS (_58_GYNGFG63^75,114^), and that of TDP-43^76^ (_58_NFGAFS_63_; also located in the protein LCD^50^). Therefore, by employing the HPS-Cation-*π* model coupled to our dynamical ageing algorithm, we can effectively investigate the viscoelastic behaviour of the FUS-LCD, A-LCD-hnRNPA1, and TDP-43-LCD condensates prior and post-maturation. Technical details on the ageing dynamical algorithm, the local order parameter driving structural transitions, and the structured interaction parameters of the coarse-grained model are provided in the SM.

We start by applying the SSRMI and OS methods to phase-separated condensates under bulk conditions of FUS-LCD, A-LCD-hnRNPA1, and TDP-43-LCD at *T* ~ 0.88 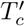 prior maturation (light brown, blue and green circles respectively in Fig. 5(a)). Please note that the critical temperature is barely affected by the maturation of the condensate, as demonstrated in our previous study^75^. As can be seen, *G*(*t*) prior maturation decays sharply in the terminal region evidencing liquid-like behaviour and full relaxation of the condensates. Moreover, we also apply the OS method to evaluate the loss and viscous moduli as a function of frequency for these condensates prior-ageing (stars in Fig. 5(b)). We find an exceptional agreement between *G′* and *G″* as a function of frequency using both OS and SSRMI techniques for FUS-LCD, A-LCD-hnRNPA1, and TDP-43-LCD condensates. Then, we activate the ageing dynamical algorithm^73,75,76^ and perform 0.4 μs simulations under bulk condensate conditions to allow inter-protein structural transitions to accumulate over time. In Fig. S2, we show the time-evolution of structural transitions driven by high-density protein fluctuations leading to inter-protein *β*-sheet domains within the condensates. For all cases, within the tested maturation time, the percentage of LARKS forming inter-protein *β*-sheet domains is higher than 75% (Fig. S2; please note that the faster dynamics of the protein model also increases the condensate maturation rate as discussed in Ref.^75^).

**FIG. 5.**
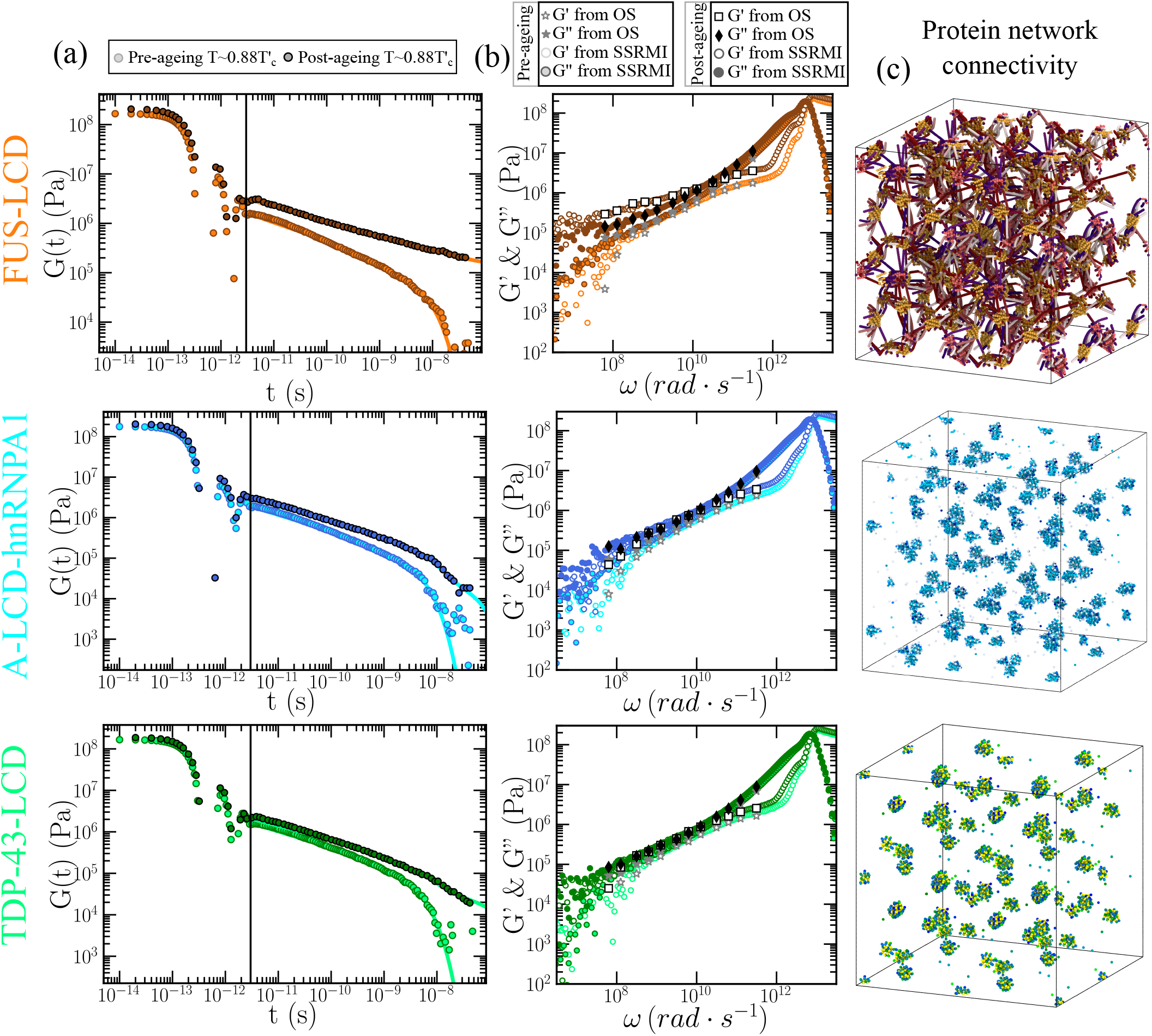
Viscoelasticity measurements and condensate network connectivity analysis of FUS-LCD, A-LCD-hnRNPA1, and TDP-43-LCD aged protein condensates. (a) Shear stress relaxation modulus of FUS-LCD (Top), A-LCD-hnRNPA1 (Middle), and TDP-43-LCD (Bottom) condensates at 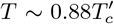 prior maturation (light colours; reference model HPS-Cation-*π*)^103,125^ and after 400 ns of maturation (dynamical ageing model^75^). (b) Elastic modulus *G′* (empty symbols) and loss modulus *G”* (filled symbols) of FUS-LCD (Top), A-LCD-hnRNPA1 (Middle), and TDP-43-LCD (Bottom) condensates from computational oscillatory shear simulations prior (stars) and after maturation (squares and diamonds). *G′* (empty circles) and *G″* (filled circles) evaluated through the Fourier transform of *G*(*t*) via the SSRMI method prior-ageing (light colours) and post-ageing (dark colours) are also included. (c) Network connectivity of aged FUS-LCD (Top), A-LCD-hnRNPA1 (Middle), and TDP-43-LCD (Bottom) condensates at 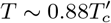 after 400 ns of maturation computed using the primitive path analysis.

We now evaluate *G*(*t*) for the FUS-LCD, A-LCD-hnRNPA1, and TDP-43-LCD aged condensates after 0.4 μs of maturation time (Fig. 5(a); darker circles). Remarkably, we find that for all condensates the observed decay in *G*(*t*) prior maturation is no longer present (light circles in Fig. 5(a)). Irrespective of the protein, ageing increases significantly the values of the shear stress relaxation modulus; hence, suggesting a much higher viscosity for aged condensates than for their pre-aged counterparts. Nevertheless, when looking more closely at the time-dependent behaviour of *G*(*t*), we observe significantly distinct curves for the different protein LCDs. While for both A-LCD-hnRNPA1 and TDP-43-LCD the continuous decay of *G*(*t*) suggests that aged condensates will present liquid-like behaviour at very long timescales (high-viscous liquids), in FUS-LCD *G*(*t*) falls into a persistent plateau with no hints of decaying at comparable timescales, and yielding infinite viscosity values (i.e., non-diffusive behaviour) characteristic of a gel-like state as recently experimentally reported for FUS^30^ and FUS-LCD^116^ condensates. The fundamental difference for FUS-LCD condensates exhibiting gelation upon condensate maturation is the presence of three separate LARKS along its sequence. At least two multivalent or three monovalent anchoring points per molecule are necessary for a system to completely gelate^117^. Thus, the strong gel-like behaviour exhibited by FUS-LCD condensates is not expected to occur in A-LCD-hnRNPA1 or TDP-43-LCD with only a single LARKS since, strictly speaking, the intermolecular network of *β*-sheets would not be able to fully percolate unless another anchoring domain along the sequence could also establish strengthened inter-protein binding (i.e., due to a sequence mutation or a post-translational modification^50^). However, according to our *G*(*t*) results shown in Fig. 5(a), both A-LCD-hnRNPA1 and TDP-43-LCD condensates may exhibit very high-viscous behaviour after maturation.

In Fig. 5(b), we also plot *G′* and *G″* post-ageing for FUS-LCD, A-LCD-hnRNPA1 and TDP-43-LCD condensates evaluated through both OS and SSRMI methods. In agreement with the results shown in Fig. 5(a), we find that for FUS-LCD condensates, *G′* upon maturation is higher than *G″*, thus indicating gel-like behaviour. In contrast, in matured A-LCD-hnRNPA1 and TDP-43-LCD condensates, the viscous modulus is higher than the elastic one, hence confirming the high-viscous liquid-like behaviour pinpointed from *G*(*t*) calculations (Fig. 5(a)) for these condensates.

To further characterise the structure and topology of the aged condensates in terms of the *β*-sheet intermolecular network, we apply a modification of the primitive path analysis (PPA) algorithm^181,182^. In our PPA calculations, we consider the *β*-sheet binding fixed in space, the bond interaction is modified to have an equilibrium bond length of 0 nm, and the intra-molecular excluded volume is set to zero. The algorithm then minimises the contour length of the protein strands that connect the different LARKS regions while preserving the topology of the underlying network. Furthermore, we replicate the system in all directions of space to better visualise the extension of the network connectivity beyond the periodic boundary conditions of the simulation box. At the end of the minimisation, this method allows the visualisation of the network connectivity generated by inter-protein *β*-sheet clusters (Fig. 5(c)). For FUS-LCD matured condensates (Fig. 5(c); Top panel), we find an elastically percolated network of protein strands that contributes to the formation of a rubbery plateau in *G*(*t*) (as shown in Fig. 5(a); Top panel). This *β*-sheet percolated network also explains the higher value of *G′* respect to *G″* upon maturation (Fig. 5(b); Top panel). On the other hand, in mature A-LCD-hnRNPA1 and TDP-43-LCD condensates, proteins form isolated *β*-sheet clusters (Fig. 5(c); Mid and Bottom panel respectively). These results from the PPA analysis are also in agreement with *G*(*t*) decaying to zero at much longer timescales (i.e., showing a higher viscosity but not a rubbery plateau as FUS-LCD), and with the viscous modulus being higher than the elastic one (Figs. 5(a) and 5(b); Middle and Bottom Panel).

Our results from Fig. 5 are fully consistent with recent experimental observations of FUS-LCD condensates where an increase in the *β*-sheet content has been associated with protein dynamical arrest within phase-separated condensates^116^. Progressive kinetic arrest through the emergence of long-lived intermolecular interactions giving rise to *β*-sheet percolated networks (Fig. 5(c)) is also consistent with the experimental observation of reversible hydrogels in LARKS-containing RNA-binding proteins after maturation (such as TDP-43 or FUS^49,50,114^) that can be dissolved with heat, and where a high percentage of *β*-sheet content has been found^116^. Furthermore, our results help to explain the recognised asphericity of aged condensates^30,183^ and the emergence of irregular morphologies caused by non-ergodic droplet coalescence^74,116^reported for LCD-containing proteins such as hnRNPA1^12^, FUS^30^, TDP-43^184^, or NUP-98^175^. Remarkably, the progressive kinetic arrest of proteins within droplets in FUS (full-sequence) in combination with a severe imbalance in the intermolecular forces has been shown to drive single-component condensates to display multiphase architectures upon maturation^73,185^ or upon phosphorylation^186^.

## III. DISCUSSION AND CONCLUSIONS

In this work, we have applied different computational techniques to characterise the viscoelastic behaviour of biomolecular condensates formed by proteins and RNA, and modelled through coarse-grained potentials of different resolution. First, by means of a simple coarse-grained model for studying IDP phase separation^100^, we have tested the validity, accuracy, and computational performance of three numerical methods to evaluate the viscosity of condensates. These methods are: the shear stress relaxation modulus integration (SSRMI)^117,118^, the oscillatory shear (OS) technique^119,120^, and the bead tracking (BT) method^30,121–123^. In Fig. 6, we summarise their different advantages and drawbacks in terms of precision, required simulation time, system size, and major computational implementation requirements. Importantly, we find that the SSRMI method is the most accurate approach to compute *η* since it does not require the extrapolation of *G”/ω* to the limit of *ω →* 0 (where the stress response signal is weak, as in the OS method), or the need of extremely long simulation timescales to avoid probe bead size-dependence (Fig. 1(b)). On the other hand, the OS approach possesses the advantage that the required simulation timescale to obtain reasonable estimates of *η* is approximately 4 and 6 times lower than the SSRMI and BT methods, respectively. Nevertheless, it relies on the implementation of a sophisticated shear deformation algorithm to perform oscillatory shear (Fig. 6). In terms of system size, while the three methods require a reasonable amount of protein replicas to avoid finite-size effects (due to protein self-interactions through the periodic boundary conditions), we note that the BT method may still demand even larger system sizes for cases where the probe radius is of the order or greater than the protein radius of gyration. However, the key advantage of this method is the simplicity of its implementation, which only requires the insertion of a probe bead with a hard sphere-like interaction with the surrounding media and the calculation of its mean square displacement within the diffusive regime. Therefore, despite the fact that each technique has its own pros and cons (Fig. 6), the SSRMI method presents the highest overall performance in terms of accuracy, implementation, and reasonable computational performance.

**FIG. 6.**
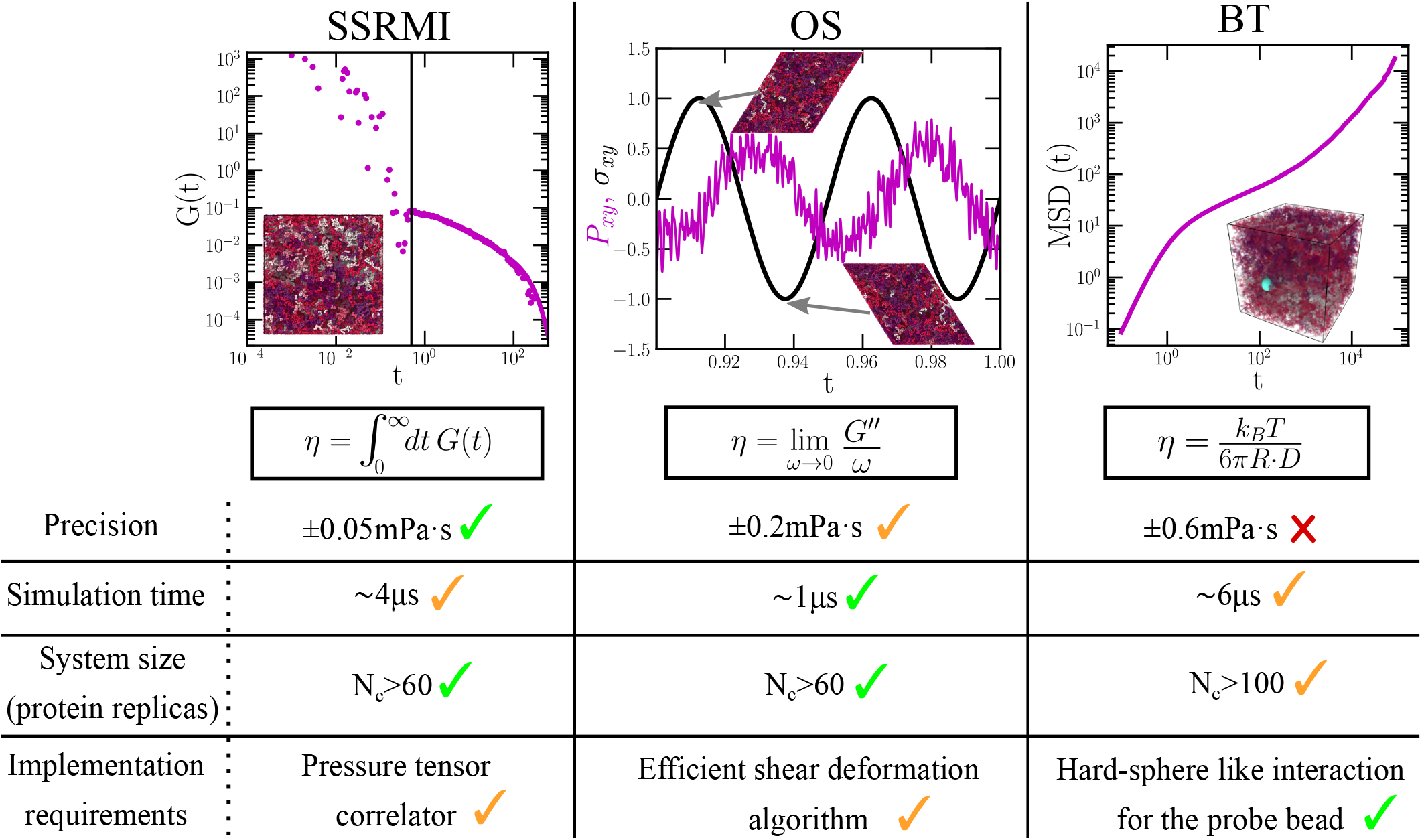
Comparison of the three different employed computational techniques to evaluate viscosity in phase-separated condensates. In the top panels we show: (Left) The decay over time of the shear stress relaxation modulus for computing *η* through the SSRMI method. (Middle) Applied shear deformation (*σ_xy_*; black curve) and stress response (*P_xy_*; purple curve) as a function of time evaluated through the oscillatory shear method. (Right) Mean square displacement of the probe bead (blue particle in the inset) to determine *η* through the Stokes-Einstein relation. Importantly, we note that SSRMI and OS calculations do not depend on the system size as long as protein self-interactions are avoided through the periodic boundary conditions whereas in BT simulations such condition might not be enough to prevent finite system-size effects in cases where the probe bead radius is greater than the protein radius of gyration. The specified data below applies for IDPs as those studied in this work.

Then, after having tested these different approaches for a simple model of IDP LLPS, we have applied the SSRMI and OS techniques for determining the condensate viscosity of a set of 7 different IDPs and 5 peptide/RNA complex coacervates using a sequence-dependent high-resolution coarse-grained model^102,125,126^ (Figs. 2 and 3). We find a reasonable agreement in the predicted viscosity between both techniques for all these systems (Figs. 3(b) and 3(c)) despite the weak stress signal of the OS method at low frequencies slightly hampers the calculation of *η* (especially for systems with low density and long chains; i.e., polyR100/polyU100). Such agreement between both techniques can be especially noticed when plotting *G′* and *G″ vs.* a wide range of frequencies for all systems, as shown in Fig. 3(a).

We also identify through our simulations a clear correlation of condensate viscosity with IDP/RNA length, molecular weight, and the number of LLPS stickers across the protein sequence (Fig. 4) when we compare all the systems at the same relative temperature with respect to the critical one. Importantly, within the stickers and spacers framework^93,167,168^, we find the best correlation when considering tyrosine, phenylalanine, and arginine as LLPS stickers^127^ than when only including aromatic residues^187^. On the contrary, when performing our calculations at constant *T* instead of 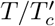, we find that IDPs and complex coacervates with higher critical temperature display higher viscosity. Similarly, higher condensate densities correlate with higher viscosities when comparing at a fixed temperature (Fig. 4(f)). Furthermore, since our simulations effectively describe the protein phase behaviour at physiological salt concentration (where electrostatic interactions are known to play a key role in sustaining LLPS^169^), we also test a possible correlation between condensate viscosity and the abundance of pairs of charged residues of distinct sign. However, a very mild trend, if any, is observed for the studied IPPs and complex coacervates as a function of the number of residue pairs with opposite charge (Fig. 4(d)).

Lastly, we have investigated by means of the SSRMI and OS methods the progressive maturation through *β*-sheet accumulation of three of the most relevant protein low-complexity domains related to the onset of ALS and FTD diseases^33,36^: A-LCD-hnRNPA^113,114,128^, FUS-LCD^49^, and TDP-43-LCD^50,129^. We find that both techniques predict the transition from condensate liquid-like behaviour to partially (A-LCD-hnRNPA1 and TDP-43-LCD) or fully (FUS-LCD) kinetically trapped states once an intermolecular *β*-sheet network has grown through the condensate, thus hindering protein self-diffusion at moderate timescales. Remarkably, the (experimentally reported^116^) emergence of gel-like behaviour in FUS-LCD condensates—due to an increase in the *β*-sheet content— can be straightforwardly identified through the OS and SSRMI methods by the higher values of *G′* with respect to *G”* at moderately low frequencies (Fig. 5(b); Top panel). Moreover, the behaviour of *G*(*t*) falling into a persistent plateau (Fig. 5(a); Top panel) corroborates the gel-like behaviour of aged FUS-LCD condensates with respect to matured A-LCD-hnRNPA1 and TDP-43-LCD droplets still presenting high-viscous liquid behaviour with much longer relaxation timescales than their pre-aged counterparts (Fig. 5(a); Middle and Bottom panel respectively). These results are also confirmed by PPA calculations revealing the underlying inter-protein *β*-sheet network emerged upon maturation (Fig. 5(c).) Taken together, our study provides a compilation of modelling rheological techniques to assess the viscoelastic properties of biomolecular condensates, which often extend far beyond those of low-viscous liquids.

## IV. ACKNOWLEDGEMENTS

This project has received funding from the Oppenheimer Research Fellowship of the University of Cambridge. A. T. is funded by Universidad Politécnica de Madrid (PhD fellowship ‘programa propio UPM’) and the Oppenheimer Fellowship. J. R. acknowledges funding from the Spanish Ministry of Economy and Competitivity (PID2019-105898GA-C22) and the Madrid Government (Comunidad de Madrid-Spain) under the Multiannual Agreement with Universidad Politécnica de Madrid in the line Excellence Programme for University Professors, in the context of the V PRICIT (Regional Programme of Research and Technological Innovation). J. R. E. also acknowledges funding from the Roger Ekins Research Fellowship of Emmanuel College. R.C.-G. acknowledges funding from the European Research Council (ERC) under the European Union Horizon 2020 research and innovation programme (grant agreement 803326). This work has been performed using resources provided by the Cambridge Tier-2 system operated by the University of Cambridge Research Computing Service (http://www.hpc.cam.ac.uk) funded by EPSRC Tier-2 capital grant EP/P020259/1. The authors gratefully acknowledge the Universidad Politécnica de Madrid (www.upm.es) for also providing computing resources on Magerit Supercomputer.

