## Supplementary Material for "Time-dependent material properties of ageing biomolecular condensates from different viscoelasticity measurements in molecular dynamics simulations"

Andrés R. Tejedor

*Maxwell Centre, Cavendish Laboratory, Department of Physics, University of Cambridge, J J Thomson Avenue, Cambridge CB3 0HE, United Kingdom. and Department of Chemical Engineering, Universidad Politécnica de Madrid, José Gutiérrez Abascal 2, 28006, Madrid, Spain*

Rosana Collepardo-Guevara

*Yusuf Hamied Department of Chemistry, University of Cambridge, Lensfield Road, Cambridge CB2 1EW, United Kingdom. and Department of Genetics, University of Cambridge, Downing Site, Cambridge CB2 3EH, United Kingdom.*

Jorge Ramírez*

*Department of Chemical Engineering, Universidad Politécnica de Madrid, José Gutiérrez Abascal 2, 28006, Madrid, Spain*

Jorge R. Espinosa†

*Maxwell Centre, Cavendish Laboratory, Department of Physics, University of Cambridge, J J Thomson Avenue, Cambridge CB3 0HE, United Kingdom.*

(Dated: December 7, 2022)

### SI. COMPUTATIONAL MODELS

#### A. A simple coarse-grained model for IDP phase separation

We use an easy coarse-grained (CG) model for intrinsically disorder protein (IDP) phase-separation to test the different computational methods to measure viscosity. The model employed is parameterized as in Ref. [1]. Our CG IDPs consists of fully flexible linear chains comprised of $N = 50$ bonded beads in which non-bonded beads interact through a Lennard-Jones potential:

$$U_{LJ}(r) = 4\epsilon \left[ \left(\frac{\sigma}{r}\right)^1 2 - \left(\frac{\sigma}{r}\right)^6 \right], \tag{S1}$$

where $r$ is the distance between two beads, $\epsilon$ is the depth of the energy well, and $\sigma$ sets the bead molecular diameter (i.e., excluded volume). For computational efficiency, the pair force computed from the gradient of $U_{LJ}$ is truncated to zero at a cut-off distance $r_c = 3\sigma$, so that non-bonded forces act only between pairs of particles with $r < r_c$. Bonded beads within a given IDP are connected by a harmonic potential:

$$U_B(r) = \frac{k}{2}(r - \sigma)^2 \tag{S2}$$

where a stiff spring constant of $k = 75000\epsilon/\sigma^2$ is used.

Typical system sizes in our IDP coarse-grained simulations contain 275 replicas. We perform Direct Coexistence simulations [2] (in the NVT ensemble) to compute (and reproduce from Ref. [1]) the phase diagram of the IDPs using the LAMMPS Molecular Dynamics package [3]. For the DC simulations we use a box with $L_x = L_y = 4L_z$ (being $L_x = 30\sigma$). The system is quenched to a temperature below the critical point, and vapor–liquid equilibrium was established over a period of $1e4\tau$. Production quality simulation data were collected over $5e4\tau$, using a Langevin thermostat [4] with a relaxation time of $5\tau$ to keep the temperature of the system constant. An integration timestep of $0.001\tau$ for the equations of motions is used.

We have estimated the viscosity of the IDP condensates using the three different methods in density-bulk equilibrium conditions (i.e., running NVT simulations of cubic boxes to

*
†

reproduce the liquid-phase conditions at the different studied temperatures). The typical length of each simulation required to compute the viscosity through SSRMI calculations is approximately $\sim 1e3\tau$. For the OS method, we apply a periodic shear deformation in the $xy$ direction, using the 'fix deform' command of the LAMMPS package, with the option 'wiggle', ranging frequencies from $\omega \sim 6 \cdot 10^{-3}\tau^{-1}$ to $\omega \sim 1\tau^{-1}$, so that the thermostat relaxation time is shorter than the deformation period, and the system is not overheated. For all frequencies, we have produced results for ∼20 periods. For the BT method, we insert a probe bead in the IDP condensate. Probe particles are modelled with an Ashbaugh-Hatch potential [5] of the following form:

$$U_{AH}(r) = \begin{cases} U_{LJ} + (1-\lambda)\epsilon, & \text{if } r \leq 2^{1/6}\sigma \\ \lambda U_{LJ}, & \text{otherwise} \end{cases}, \tag{S3}$$

where $U_{LJ}$ refers to the standard Lennard-Jones potential as described in Eq. S1, and $\lambda$ is set to zero in order to establish a purely repulsive interaction between the probe bead and the IDP chains. The mass of the probe bead is set to $m = 1$ (in reduced units; see main text), the potential energy depth is set to $\epsilon = 4$ to avoid interpenetration of the chains and the boundary of the probe bead, and the cut-off distance for $U_{AH}$ interactions is 3 times the probe bead molecular diameter. Equilibration is carried out through an energy minimisation algorithm using 2000 evaluations with a tolerance of $10^{-4}$ and $10^{-6}$ in energy and force respectively. Production runs are of the order of $\sim 10^4\tau$ or below, depending on the temperature and timescale needed so that the mean-squared displacement of the probe bead converges to the Fickian regime.

#### B. HPS-Cation−$\pi$ model

In order to simulate the different studied proteins and complex coacervates (proteins and RNA), we employ the LAMMPS Molecular Dynamics simulation package [3] with the recent reparameterization by Das *et al.* [6] of the chemically-accurate coarse-grained HPS protein model proposed by Dignon *et al.* [7]. For RNA, we use the HPS-compatible CG model proposed by Regy *et al.* [8]. The coarse-grained model resolution, both for proteins and RNA, is of one bead per amino acid and nucleotide. In the model, intrinsically disordered proteins and RNA strands are considered as fully flexible polymers. The potential energy of the HPS-Cation$-\pi$ force field is given by:

$$E = E_{\text{Bonds}} + E_{\text{Electrostatic}} + E_{\text{Hydrophobic}} + E_{\text{Cation}-\pi}, \tag{S4}$$

where $E_{\text{Hydrophobic}}$, $E_{\text{Cation}-\pi}$ and $E_{\text{Electrostatic}}$ interactions are only applied between non-bonded beads (i.e., residues or nucleotides), and $E_{\text{Bonds}}$ between subsequent beads directly bonded to each other. Bonded interactions between subsequent amino acid protein beads or consecutive RNA nucleotides are described by a harmonic potential:

$$E_{\text{Bonds}} = \sum_{\text{Protein/RNA bonds}} k(r - r_0)^2, \tag{S5}$$

where the equilibrium bond length between subsequent nucleotides is $r_0 = 5.0$ Å, and $r_0 = 3.81$ Å between bonded amino acid beads. The spring constant is $k = 10\,\text{kJ/(mol\AA}^2)$. Electrostatic interactions, $E_{\text{Electrostatic}}$, among charged amino acids and RNA nucleotides are described by a Yukawa/Debye-Hückel potential of the following form:

$$E_{\text{Electrostatic}} = \sum_i \sum_{j<i} \frac{1}{4\pi D} \frac{q_i q_j}{r} e^{-r/\kappa}, \tag{S6}$$

where $q_i$ and $q_j$ represent the charges of the beads $i$ and $j$ (amino acids or nucleotides), $D = 80\varepsilon_0$ is the relative dielectric constant of water (being $\varepsilon_0$ the permittivity of vacuum), $r$ is the distance between the $i$th and $j$th beads, and $\kappa = 1$ $nm$ is the Debye screening length that mimics the implicit solvent (water and ions) at physiological salt concentration (∼150mM of NaCl) [7].

Hydrophobic interactions between different amino acids and nucleotides are built upon an hydrophobicity residue/nucleotide scale based on a statistical potential derivation from contacts in PDB structures, and implemented through the functional form of an Ashbaugh/Hatch potential (see further details on these References [5, 7–9]):

$$E_{\text{Hydrophobic}} = \sum_i \sum_{j<i} \begin{cases} 4\epsilon_{ij} \left[ \left(\frac{\sigma_{ij}}{r}\right)^{12} - \left(\frac{\sigma_{ij}}{r}\right)^{6} \right] + (1 - \lambda_{ij})\epsilon_{ij}, & r < 2^{1/6}\sigma_{ij} \\ \lambda_{ij} 4\epsilon_{ij} \left[ \left(\frac{\sigma_{ij}}{r}\right)^{12} - \left(\frac{\sigma_{ij}}{r}\right)^{6} \right], & \text{otherwise,} \end{cases} \tag{S7}$$

where $\lambda_i$ and $\lambda_j$ are parameters that account for the hydrophobicity of the $i$th and $j$th interacting particles respectively, being $\lambda_{ij} = (\lambda_i + \lambda_j)/2$. The excluded volume of the different residues/nucleotides is given by $\sigma_i$ and $\sigma_j$, where $\sigma_{ij} = (\sigma_i + \sigma_j)/2$, and $r$ is the distance between the $ij$ particles. $\epsilon_{ij}$ (0.2 kcal/mol) is a fitting parameter to reproduce the experimental single-IDP radius of gyration ($R_g$) of a protein set as shown in Ref. [7]. The specific values for each amino acid and nucleotide $\sigma$, $q$, and $\lambda$ parameters can be found in References: Dignon *et al.* [7] for proteins, and Regy *et al.* [8] for RNA.

Finally, we consider an extra term to describe cation-$\pi$ interactions (only for the following set of pairs of amino acids (c-$\pi$:{Arg-Phe, Arg-Trp, Arg-Tyr, Lys-Phe, Lys-Trp and Lys-Tyr}):

$$E_{\text{cation-}\pi} = \sum_{i \in \text{c-}\pi} \sum_{j \in \text{c-}\pi \& j<i} 4\epsilon_{ij} \left[ \left( \frac{\sigma_{ij}}{r} \right)^{12} - \left( \frac{\sigma_{ij}}{r} \right)^{6} \right], \tag{S8}$$

where $\sigma_{ij}$ is the same as in the hydrophobic interactions, and $\epsilon_{ij}$ is $\epsilon_{ij} = 3.0\mathrm{kcal\,mol}^{-1}$ for all six cation-$\pi$ pairs as proposed in Ref. [6] (Approach 1).

We run Direct Coexistence (DC) simulations [2] to compute phase diagrams in the NVT ensemble. We use a Langevin thermostat [4] with a relaxation time of 5 ps, and a timestep of 10 fs for the Verlet integration of the equations of motion.

Viscosity has been estimated using both SSRMI and OS methods independently, under bulk density conditions for all the IDPs and complex coacervates studied. The calculation of $G(t)$ has been carried out running cubic NVT simulations. The typical length of each simulation to compute viscosity is of ∼5 $\mu$s. Oscillatory shear simulations have been carried out using the 'fix deform' command of the LAMMPS package, with the option 'wiggle', ranging frequencies below $\omega \sim 0.1\,ns^{-1}$. LARKS disorder-to-order transitions are implemented through our dynamical ageing model (as discussed in Section SV) in bulk phases at condensate coexistence densities (extracted from DC simulations and our previous works [10, 11]) running NVT simulations.

### SII. PHASE DIAGRAM CALCULATIONS

The phase diagrams of the different studied systems (Figs. 1a, 2a, and 2b of the main text) have been evaluated by means of Direct Coexistence simulations [2]. The two coexisting phases are introduced in the same simulation box, and the system is equilibrated at a given temperature. From the density profile, computed along the long axis of the simulation box, we evaluate the density of the condensed and diluted phases. The critical point of the phase diagrams is estimated using the universal scaling law of coexistence densities near a critical point [12], and the law of rectilinear diameters [13]:

$$\big(\rho_l(T) - \rho_v(T)\big)^{3.06} = d\bigg(1 - \frac{T}{T_c}\bigg), \tag{S9}$$

and

$$(\rho_l(T) + \rho_v(T))/2 = \rho_c + s_2(T_c - T), \tag{S10}$$

where $\rho_v$ and $\rho_l$ are the coexisting densities of the diluted and condensed phases, respectively, $T_c$ and $\rho_c$ are the critical temperature and density, respectively, and $d$ and $s_2$ are fitting parameters.

### SIII. PROTEIN SEQUENCES

#### $\alpha$-synuclein

MDVFMKGLSKAKEGVVAAAEKTKQGVAEAAGKTKEGVLYVGSKTKEGVVHGVATVAEKTKEQ
VTNVGGAVVTGVTAVAQKTVEGAGSIAAATGFVKKDQLGKNEEGAPQEGILEDMPVDPDNEAYE
MPSEEGYQDYEPEA

#### Ddx4

MGDEDWEAEINPHMSSYVPIFEKDRYSGENGDNFNRTPASSSEMDDGPSRRDHFMKSGFASGRNF
GNRDAGECNKRDNTSTMGGFGVGKSFGNRGFSNSRFEDGDSSGFWRESSNDCEDNPTRNRGFSK
RGGYRDGNNSEASGPYRRGGRGSFRGCRGGFGLGSPNNDLDPDECMQRTGGLFGSRRPVLSGTG
NGDTSQSRSGSGSERGGYKGLNEEVITGSGKNSWKSEAEGGES

#### LAF-1-RGG

MESNQSNNGGSGNAALNRGGRYVPPHLRGGDGGAAAAASAGGDDRRGGAGGGGYRRGGGNSGG
GGGGGYDRGYNDNRDDRDNRGGSGGYGRDRNYEDRGYNGGGGGGGNRGYNNNRGGGGGGYN
RQDRGDGGSSNFSRGGYNNRDEGSDNRGSGRSYNNDRRDNGGDGLEHHHHHH

#### Tau K18

MQTAPVPMPDLKNVKSKIGSTENLKHQPGGGKVQIINKKLDLSNVQSKCGSKDNIKHVPGGGSVQI
VYKPVDLSKVTSKCGSLGNIHHKPGGGQVEVKSEKLDFKDRVQSKIGSLDNITHVPGGGNKKIE

#### TDP-43-LCD

GRFGGNPGGFGNQGGFGNSRGGGAGLGNNQGSNMGGGMNFGAFSINPAMMAAAQAALQSSWG
MMGMLASQQNQSGPSGNNQNQGNMQREPNQAFGSGNNSYSGSNSGAAIGWGSASNAGSGSGFNG

GFGSSMDSKSSGWGM

**FUS-LCD**

MASNDYTQQATQSYGAYPTQPGQGYSQQSSQPYGQQSYSGYSQSTDTSGYGQSSYSSYGQSQNTG
YGTQSTPQGYGSTGGYGSSQSSQSSYGQQSSYPGYGQQPAPSSTSGSYGSSSQSSSYGQPQSGSYSQ
QPSYGGQQQSYGQQQSYNPPQGYGQQNQYNS

**A-LCD-hnRNPA1**

MASASSSQRGRSGSGNFGGGRGGGFGGNDNFGRGGNFSGRGGFGGSRGGGGYGGSGDGYNGFG
NDGSNFGGGGSYNDFGNYNNQSSNFGPMKGGNFGGRSSGPYGGGGQYFAKPRNQGGYGGSSSSSS
YGSGRRF

**$PR_{25}$**

PRPRPRPRPRPRPRPPRPRPRPRPRPRPRPRPRPRPRPRPRPRPRPPRPRPR

### SIV. ADDITIONAL TECHNICAL DETAILS FOR THE VISCOSITY CALCULATIONS

#### A. Shear Stress Relaxation Modulus Integration (SSRMI)

From NVT simulations, we compute the viscosity of the condensates using both the simple IDP model [1] and the HPS-Cation$-\pi$ force field [7, 14]. In an isotropic system, we can accurately calculate the shear relaxation modulus $G(t)$ using all components of the pressure tensor ($\sigma_{\alpha\beta}$) as shown in Ref. [15]:

$$\begin{aligned} G(t) = &\frac{V}{5k_BT}\left[\langle\sigma_{xy}(0)\sigma_{xy}(t)\rangle + \langle\sigma_{xz}(0)\sigma_{xz}(t)\rangle + \langle\sigma_{yz}(0)\sigma_{yz}(t)\rangle\right] \\ &+ \frac{V}{30k_BT}\left[\langle N_{xy}(0)N_{xy}(t)\rangle + \langle N_{xz}(0)N_{xz}(t)\rangle + \langle N_{yz}(0)N_{yz}(t)\rangle\right], \end{aligned} \quad \text{(S11)}$$

where $N_{\alpha\beta} = \sigma_{\alpha\alpha} - \sigma_{\beta\beta}$ is the first normal stress difference. This correlation can be easily and efficiently computed using the 'compute ave/correlate/long' in the 'USER-MISC' package of LAMMPS, where a multiple-$\tau$ correlator is implemented[15]. The viscosity can then be obtained by calculating the integral of $G(t)$ from $t = 0$ to $t = \infty$. The initial regime, up to $t = t_0$, accounting for intramolecular interactions and bond/angles relaxation, is integrated numerically using the trapezoidal rule. The terminal region, from $t = t_0$ to $\infty$, corresponding to slower relaxation modes (as conformational relaxation, intermolecular interactions, or self-diffusion within a crowded environment), is treated by fitting the curve of $G(t)$ to the sum of 4 to 6 Maxwell modes, depending on the span and noise of $G(t)$. The series of Maxwell modes ($G_i \exp(-t/\tau_i)$) are equidistant in logarithmic time [16], and the fit is carried out with the help of the open-source RepTate software [17]. Finally, viscosity is obtained by adding the two terms:

$$\eta = \eta(t_0) + \sum_{i=0}^{m} \tau_i G_i \exp\Big( - t_0/\tau_i \Big) \tag{S12}$$

where $\eta(t_0)$ corresponds to the computed term for short time scales, $t_0$ is the time that separates both regimes, $m$ is the number of Maxwell modes employed in the fit, and $\tau_i$ and $G_i$ are the fitting parameters of the Maxwell modes. Simulations of 3 to 5 $\mu$s, depending on the system, are needed to observe a clear terminal decay in $G(t)$, characteristic of liquids. Storage ($G'$) and loss ($G''$) moduli can be directly calculated by applying the Fourier transform to $G(t)$, so that the complex modulus $G^*(\omega)$ is obtained, and $G'$ corresponds to the real part and $G''$ to the complex part of $G^*(\omega)$. In our case, RepTate software [17] provides the calculation of the Fourier transform of $G(t)$ by default.

#### B. Oscillatory shear (OS) technique

This method relies on applying an oscillatory shear deformation to the simulation box, emulating the small-angle oscillatory shear (SAOS) experiment which is typical of bulk rheology. In the LAMMPS package, this is implemented by means of the 'fix deform' command with the option 'wiggle', which requires a triclinic instead of an orthogonal box, and simulations are carried out in the NVT ensemble to keep the temperature of the system constant. Details about the calculation of $G'$ and $G''$ through the OS method are provided in the main text, however, here we provide some additional technical details.

Similar to the SAOS experiments, before performing the frequency sweep, we must identify the linear viscoelastic (LVE) regime by doing an amplitude sweep. We select an intermediate frequency for representative systems at the temperature of study: $T = 0.96T_c$ for the simple IDP model (Fig. 1 of the main text), or $T \sim 0.88T_c'$ for condensates such as those simulated through the HPS-Cation-$\pi$ force field (i.e., Ddx4 for the studied set of IDPs, or polyR100/polyU100 for the complex coacervates). At the test frequency, we increase the deformation amplitude from $\gamma_0 = 0.05$ to $\gamma_0 = 1.5$. The frequency must be sufficiently slow (*e.g.*, 10 times slower than the relaxation of the thermostat) to avoid system overheating, but high enough to detect a mechanical response of the system at small amplitudes that clearly exceeds the thermal fluctuations of the stress. The LVE regime is identified in the region where the amplitude of the response grows linearly with $\gamma_0$, or equivalently, the regime at which $G'$ and $G''$ are independent of the amplitude of the applied deformation. In all of our tested systems, the LVE regime was identified within the entire range of amplitudes that were explored with no significant increase in temperature in any case.

To ensure that the frequency sweep is performed inside the LVE regime, we restrict the amplitude deformation to $\gamma_0 = 0.6L$. Please note that in experiments this amplitude would be considered as too large to be in the linear regime. However, our previous amplitude sweeps show that the linear approximation still holds under these conditions. In order to determine the optimal range of frequencies to perform OS simulations, we must take into account the following considerations:

- The maximum frequency must be much slower than the relaxation of the thermostat.
- The signal to noise ratio of the measured stress must be acceptable, so that of the we can perform a sinusoidal fit to the stress.
- Systems with lower densities present more difficulty in producing a significant response (due to the higher noise to signal ratio).
- Several periods of deformation must be run to improve the fit statistics.
- In order to measure the viscosity, it is important to reach for the lowest possible measurable frequency, so that $G'' \propto \omega$.

In all the systems, the above conditions were fulfilled, but in some particular cases the amplitude and frequency had to be modified. For Ddx4 and Tau K18 condensates, the regime $G'' \propto \omega$ is reached at lower frequencies, so that we also explored ($\omega \sim 1 \cdot 10^8\, s^{-1}$), increasing the deformation to $\gamma_0 = 0.9L$ to consequently increase the response amplitude. Moreover, since the complex coacervates present a relatively low density, the amplitude has been increased to $\gamma_0 = 0.9L$ for frequencies $\omega \leq 1 \cdot 10^{10}\, s^{-1}$, hence, ensuring a good fit to the response of the system, while still being inside the LVE regime.

For the systems that undergo ageing, we have used the final configuration after running NpT simulations allowing crossed-$\beta$-sheet transitions (Fig. S2). Then, fixing the density of each system to its bulk value after ageing, they have been equilibrated for 10 ns. OS simulations have been carried out in the NVT ensemble (with the dynamic ageing algorithm switched off).

Additionally, we looked into the possibility of using the OS technique in-place as the aging process progressed, which is one of the standard experimental procedures to detect gellation. Starting from a pre-aged configuration and enabling the dynamical algorithm to allow transitions, we applied the shear oscillatory deformation. As the system undergoes ageing, the mechanical response is expected to change, with an increasing elastic response. However, in our tests, the angular phase between the deformation and the shear stress response remained almost constant. This result can be somehow expected since condensates usually increase their density upon maturation [10], but in OS simulations the box volume must be kept fixed.

#### C. Bead Tracking (BT) method

The bead tracking method has been employed to estimate the condensate viscosity in our simple coarse-grained model for IDP phase-separation (Sec. SI A). In that section along with the main text, we provide the details to simulate and implement the bead tracking technique. However, the most important technical considerations can be summarized in the following points:

- The probe bead immersed in the condensate must have a steep purely repulsive interaction with the medium.
- It is recommended to use one single bead to avoid cooperative effects and bead clustering.
- The radius of the probe bead must be large enough to sample the medium but much smaller than the box size. If the medium forms some kind of network, the probe size must be larger than the characteristic mesh size.

- Simulations must be run at least one decade longer than the required time for reaching the terminal Fickian time of the probe bead.

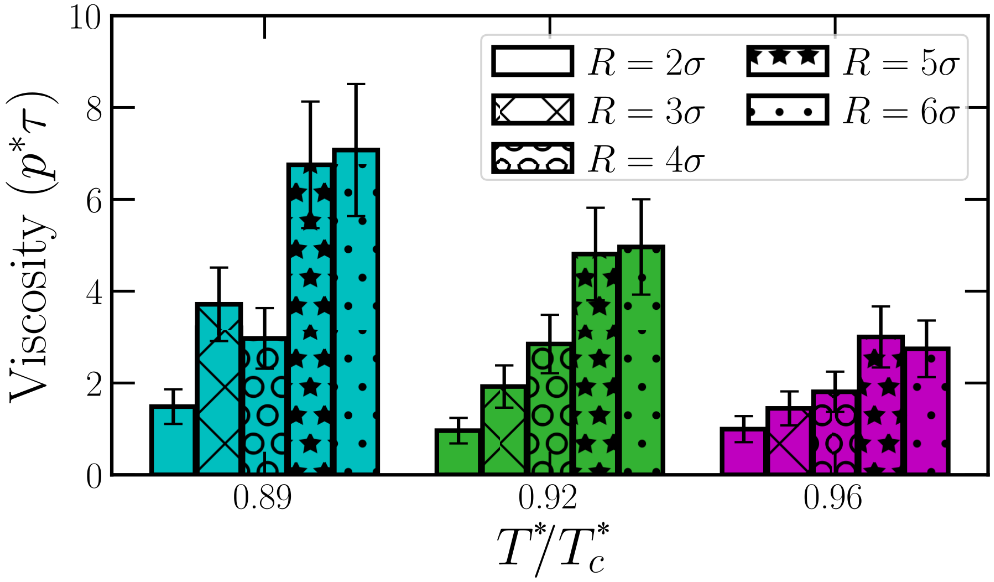


**FIG. S1:** Viscosity evaluated through BT calculations for condensates modelled through the simple IDP coarse-grained model using probe beads with different radii (as indicated in the legend), and at different temperatures. Please note that $\sigma$ refers to the bead molecular diameter of the IDP chains.

In Fig. S1 we show the viscosity estimation for IDP condensates using different probe bead radii. As can be seen, while at high temperatures all the explored radii (except that of $R = 2\sigma$) agree within the uncertainty in estimating the viscosity, at lower temperatures the beads with $R$=5 or 6 $\sigma$ do not sample the system properly. Furthermore, for the smallest considered probe particle ($R = 2\sigma$), the viscosity of the medium is underestimated since the bead size is of the order of the mesh size of the system. We note that the estimated error for $\eta$ using this technique, for all beads and temperatures, is sensibly larger than those from SSRMI and OS calculations (Fig. 1 of the main text). Instead of directly measuring the diffusion coefficient from the MSD of the bead, we could alternatively apply the Fourier transform to $g_3(t)$ and from that infer $G'$ and $G''$ (see Ref. [18] for technical details on this possible additional calculation).

### SV. DYNAMIC ALGORITHM FOR CONDENSATE AGEING SIMULATIONS

In this work, we have also employed the dynamic ageing algorithm developed in our previous studies [10, 19]. Through this algorithm, the Low-complexity Aromatic-Rich Kinked Segments (LARKS) of FUS-LCD, A-LCD-hnRNPA1, and TDP-43-LCD can undergo 'effective' structural transitions from disordered states to inter-protein $\beta$-sheets (in terms of binding energy and structure) depending on the local environment and in a time dependent manner. The LARKS sequences of these different IDPs are:

- FUS-LCD: $_{37}$SYSGYS$_{42}$, $_{54}$SYSSYGQS$_{61}$, and $_{77}$STGGYG$_{82}$
- A-LCD-hnRNPA1: $_{58}$GYNGFG$_{63}$
- TDP-43-LCD: $_{39}$NFGAFS$_{44}$

To recapitulate in our simulations such transitions, every 100 simulation timesteps, if the conditions around each of the disordered LARKS are favourable to form a cross-$\beta$-sheet motif (i.e., a high-density fluctuation), the dynamic algorithm changes the force field parameters to those given in Table S1 corresponding to inter-protein structured $\beta$-sheets. The dynamical evaluation of these conditions takes place on the central bead of each LARKS. We employ a cut-off distance criterion ($r_{cut}$ in Table S1) to evaluate the required conditions and trigger the structural $\beta$-sheet transitions. This occurs when four LARKS met within a given cut-off distance (Table S1) [10, 19, 20]. Then, the average effective interaction binding between these LARKS is replaced by that obtained from atomistic PMF simulations (see Refs. [10, 19, 20]). In this work, inter-protein $\beta$-sheet transitions are reversible for all the systems. If one of the structured LARKS separates from the crossed-$\beta$-sheet motif beyond a given distance ($r_{cut,reverse}$ in Table S1), the interaction parameters are reversed to those of the disorder residues (HPS-Cation-$\pi$ force field). After a structural transition takes place, we assign an average mass to the new (averaged) structured LARKS residues that keeps the total mass of the proteins constant. In addition, we mimic the local increase of rigidity associated with a $\beta$-sheet structural transition by introducing an angular term to the total energy (Eq. (S4)):

$$E_{\text{Angle}} = \sum_{\text{Angles}} k_{ang}(\theta - \theta_0)^2, \tag{S13}$$

where we set $\theta_0 = 180°$ and $k_{ang} = 5\ kcal\ mol^{-1}\ rad^{-2}$ for the structured LARKS of an inter-protein $\beta$-sheet. To carry out these simulations, we use the USER-REACTION [21] package of LAMMPS that allows us to change the topology and identity of the selected residues (i.e., LARKS within a LARKS high-density fluctuation) in a time-dependent and local-dependent manner.

| | A-LCD-hnRNPA1 | FUS (1) | FUS (2) | FUS(3) | TDP-43-LCD |
|---|---|---|---|---|---|
| Central amino acid | $N_{60}$ | $S_{39}$ | $S_{57}$ | $G_{79}$ | $A_{42}$ |
| $m_{ordered}$ / (g $mol^{-1}$) | 99.27 | 107.44 | 107.48 | 86.92 | 103.95 |
| $\lambda_{ordered}$ / (kcal $mol^{-1}$) | 3.2 | 2.8 | 3.5 | 2.2 | 3.5 |
| $\sigma_{ordered}$ / Å | 5.33 | 5.52 | 5.52 | 5.13 | 5.52 |
| $r_{cut}$ / Å | 8 | 7 | 7 | 7.75 | 7.75 |
| $r_{cut,reverse}$ / Å | 20 | 20 | 20 | 20 | 20 |

**TABLE S1:** Set of parameters employed for residues belonging to structured inter-peptide $\beta$-sheet motifs in HPS-Cation-$\pi$ simulations. FUS (1), FUS (2) and FUS (3) correspond to $_{37}$SYSGYS$_{42}$, $_{54}$SYSSYGQS$_{61}$ and $_{77}$STGGYG$_{82}$ sequences respectively.

The criterion that at least four peptides should be in close contact to trigger a structural disorder-to-order transition has been chosen because it is the minimal system where all different types of stacking and hydrogen bonding interactions that stabilise the $\beta$-sheet fibrillar ladder are fulfilled (please see discussion in Ref. [19]). In fact, if we consider fewer interacting peptides (e.g., only two or three), the strength of interactions among ordered LARKS would be severely lower than that reported by potential of mean force atomistic simulations [10, 19, 22] for 4 and 6 peptides forming a crossed-$\beta$-sheet, and thus, the number of transitions back to unstructured states would be high, making the ageing algorithm highly inefficient.

In Fig. S2, we show the time-evolution of inter-protein $\beta$-sheet structural transitions for FUS-LCD, TDP-43-LCD, and A-LCD-hnRNPA1 at the same relative temperature to their corresponding critical temperature ($T/T_c' \sim 0.88$). Simulations have been carried out in the NpT ensemble to allow the condensate density to relax during ageing. The resulting configurations of the condensates upon 400 ns of maturation have been then used to measure the impact of inter-protein $\beta$-sheet accumulation on the viscoelastic behaviour of aged condensates.

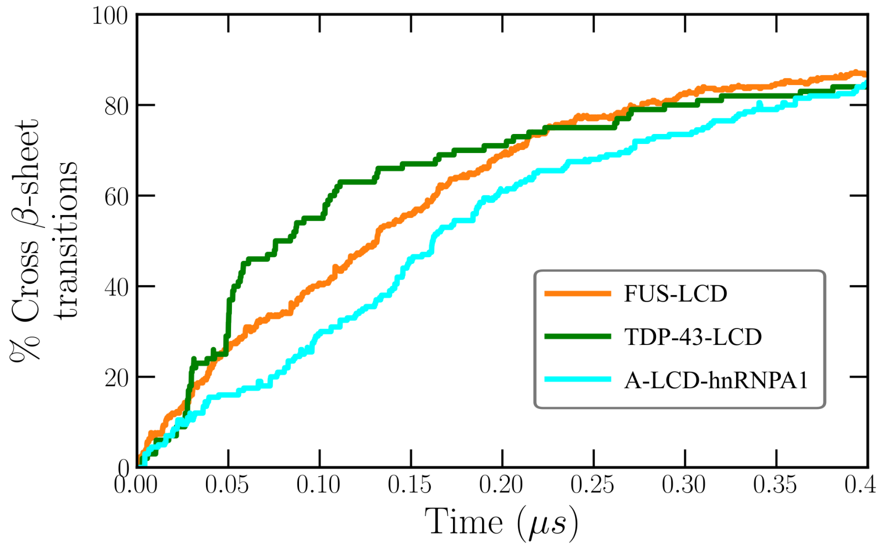


**FIG. S2:** Percentage of LARKS forming inter-peptide $\beta$-sheet transitions as a function of time in phase-separated condensates of FUS-LCD, A-LCD-hnRNPA1, and TDP-43-LCD (as indicated in the legend) at $T/T_c' \sim 0.88$ (being $T_c'$ the corresponding critical temperature of each system).

### SVI. ANALYSIS OF THE NETWORK CONNECTIVITY BY A PRIMITIVE PATH ANALYSIS

We analyse the $\beta$-sheet network connectivity of the aged condensates using the primitive path analysis (PPA) method [23] in Fig. 5c of the main text. PPA minimises the contour length of all chains in the condensate, while keeping the end monomers fixed, and thus respecting the network topology by not allowing chains to cross each other. We have employed a modification of this algorithm proposed by Tejedor *et al.* [10]. For this purpose, we first perform NpT simulations to allow the number of cross-$\beta$-sheet transitions in each of the different studied proteins to reach a steady state. Within the employed modification, we freeze the positions of the inter-protein $\beta$-sheet motifs. Then, we perform an energy minimisation in which the bond length is fixed to zero and where the excluded volume interactions are removed. Thus, the contour length of the strands between the transitioned structured LARKS is minimised while preserving the underlying network connectivity. To run the energy minimization required for the PPA method, we have used LAMMPS [3] using $1{\cdot}10^5$ iterations with a tolerance of $1{\cdot}10^{-6}$ for both energy and total force.

### SVII. ADDITIONAL SUPPORTING FIGURES

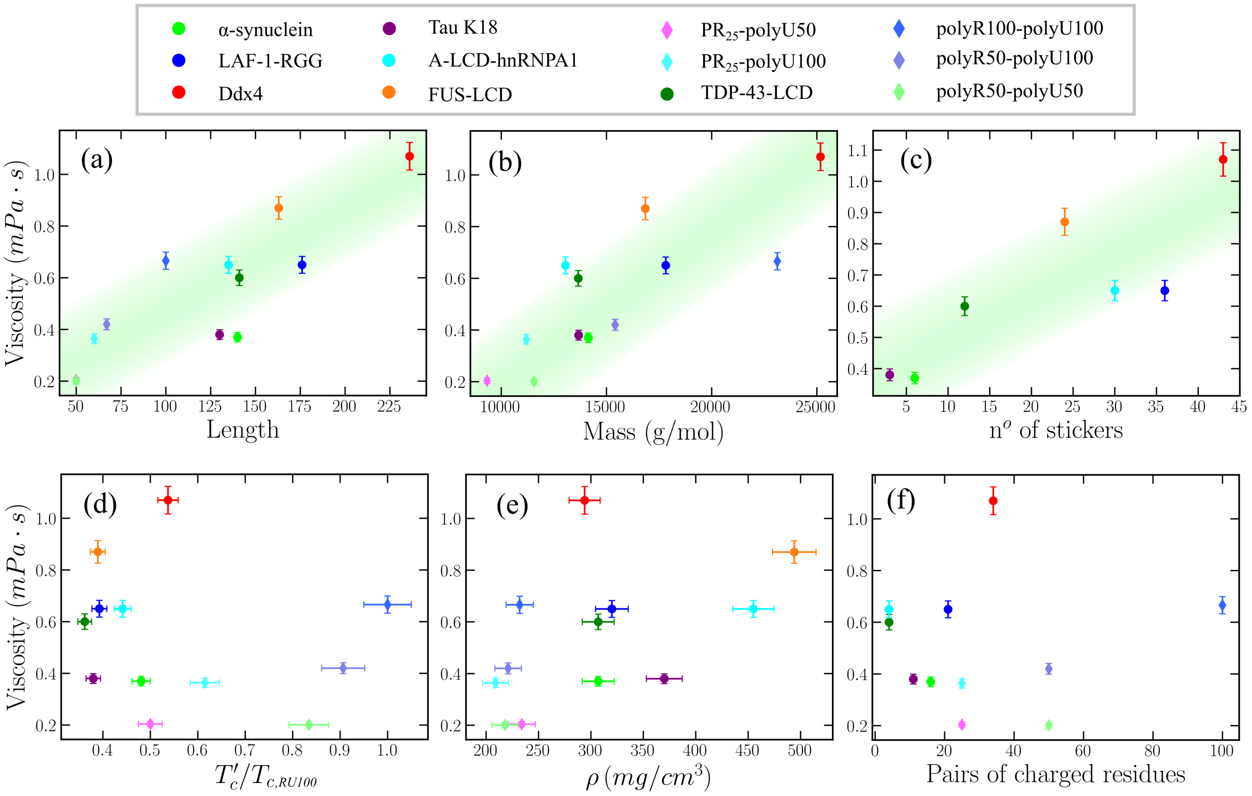


**FIG. S3:** Correlation between condensate viscosity (obtained via the SSRMI method) and chain length (a), molecular weight (b), number of stickers (Y, F and R) across the sequence (c), system critical temperature $T'_c$ (d), condensate density (e), and number of charged residue pairs of opposite charge (f) at $T \sim 0.88T'_c$ for all systems. In complex coacervates, we take the average chain length and molecular weight of the two cognate molecules. In panel (d), we note that each critical temperature has been renormalized by the highest critical temperature of the studied set ($T_{c,RU100} \equiv T_{c,\mathrm{polyR100/polyU100}}$, which corresponds to that of polyR100/polyU100). Since the complex coacervates by construction do not contain aromatic residues, their results have not been considered for the correlation shown in panel (c).

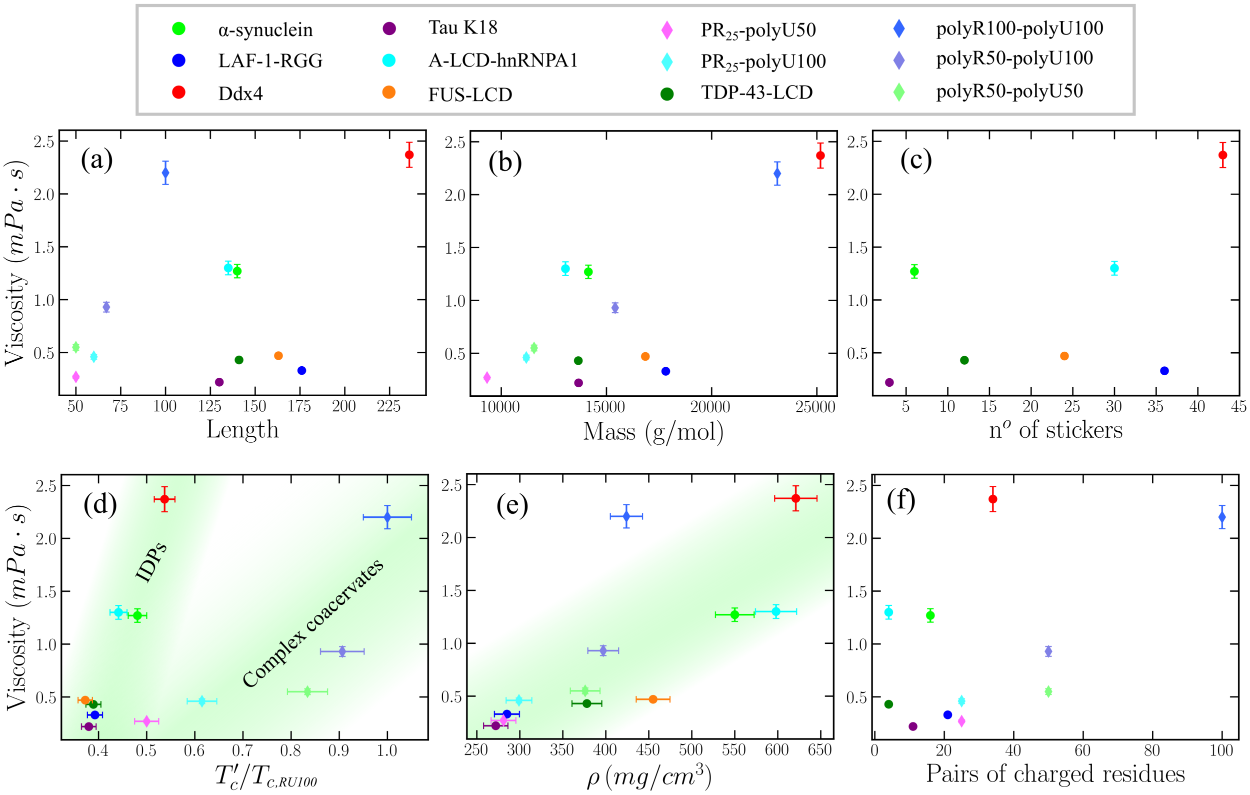


**FIG. S4:** Correlation between condensate viscosity (obtained via the SSRMI method) and chain length (a), molecular weight (b), number of stickers (Y, F and R) across the sequence (c), system critical temperature $T_c'$ (d), condensate density (e), and number of charged residue pairs of opposite charge (f) at $T = 340K$ for all systems. In complex coacervates, we take the average chain length and molecular weight of the two cognate molecules. In panel (d), we note that each critical temperature has been renormalized by the highest critical temperature of the studied set ($T_{c,RU100}$, which corresponds to that of polyR100/polyU100). Since the complex coacervates by construction do not contain aromatic residues, their results have not been considered for the correlation shown in panel (c).

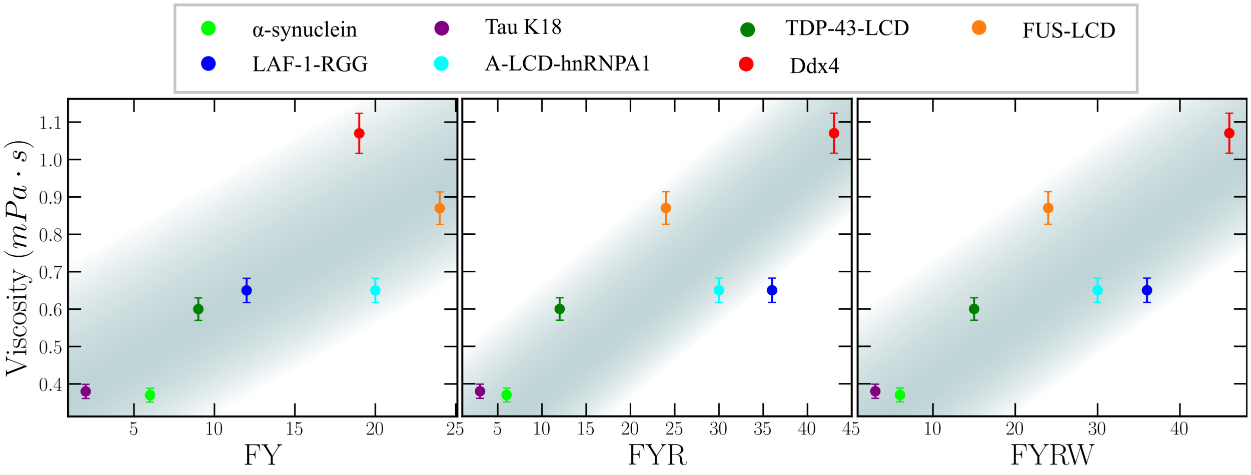


**FIG. S5:** Correlation between condensate viscosity and number of stickers along the sequence, defining stickers as F and Y (left); F, Y, and R (center, and main text; Fig. 4c); and F, Y, R, and W (right), at $T \sim 0.88T_c'$ for all systems. The different colours correspond to the different studied IDPs described in the legend.

---

[6] S. Das, Y.-H. Lin, R. M. Vernon, J. D. Forman-Kay, and H. S. Chan, "Comparative roles of charge, $\pi$, and hydrophobic interactions in sequence-dependent phase separation of intrinsically disordered proteins," *Proceedings of the National Academy of Sciences*, vol. 117, no. 46,

pp. 28795–28805, 2020.

[12] J. S. Rowlinson and B. Widom, *Molecular theory of capillarity.* Courier Corporation, 2013.

[20] S. Blazquez, I. Sanchez-Burgos, J. Ramírez, T. Higginbotham, M. M. Conde, R. Collepardo-Guevara, A. R. Tejedor, and J. R. Espinosa, “Reordering of aromatic-rich segments in fus inhibits ageing of fus-rna condensates,” *bioRxiv*, 2022.

[21] J. R. Gissinger, B. D. Jensen, and K. E. Wise, “Modeling chemical reactions in classical molecular dynamics simulations,” *Polymer*, vol. 128, pp. 211–217, 2017.

[22] A. Garaizar, J. R. Espinosa, J. A. Joseph, and R. Collepardo-Guevara, “Kinetic interplay between droplet maturation and coalescence modulates shape of aged protein condensates,” *Scientific reports*, vol. 12, no. 1, pp. 1–13, 2022.

[23] S. K. Sukumaran, G. S. Grest, K. Kremer, and R. Everaers, “Identifying the primitive path mesh in entangled polymer liquids,” *Journal of Polymer Science Part B: Polymer Physics*, vol. 43, no. 8, pp. 917–933, 2005.
